# Sodium Butyrate Rescues Neurodevelopmental Deficits Following Perinatal Methadone Exposure

**DOI:** 10.1101/2025.06.26.661851

**Authors:** Isobel A. R. Williams, Josie van Dorst, Sarah-Jane Leigh, Sarah. J. Baracz, B. L. D. Uthpala Pushpakumara, Abigail Marcus, Delyse McCaffrey, Adam K. Walker, Chee Y. Ooi, Meredith C. Ward, Ju-Lee Oei, Kelly J. Clemens

**Author notes:** Corresponding author: Kelly Clemens, PhD, School of Psychology, University of New South Wales, Kensington, NSW 2052.

## Abstract

Prenatal opioid exposure (POE) induces long-term neurodevelopmental, behavioral and cognitive deficits for which no targeted treatments are available. The mechanisms underlying POE deficits are poorly understood, but have been linked to a range of central, peripheral, and enteric nervous system changes. Emerging evidence indicates that maternal microbiota changes may also contribute to these long-term deficits in offspring. Here we test the efficacy of the short-chain fatty acid sodium butyrate (NaB) to mitigate POE-induced deficits in a rat model. Both methadone and sodium butyrate treatments altered dam microbiota composition and function: notably methadone disrupted dam gene expression of microbial enzymes critical for butyric acid production and reduced faecal butyric acid levels. In postnatal day 9 pups, methadone increased gut barrier permeability that was reversed with NaB, and enzymatic disruptions were observed in pups at postnatal day 21 that resolved in adulthood. POE induced anxiety-like behavior in adolescence, and adult deficits in working spatial memory and attentional processing that were partially rescued in rats that had received prenatal NaB. POE was associated with decreased myelination in the hippocampus, and this was partially reversed by NaB. Together these results highlight for the first time the link between the gut-brain axis in animal models of POE. Furthermore, they provide the first indication in a rat model of NaB as a simple yet effective treatment to significantly improve the outcomes of children born with POE.

## Introduction

Opioid use is at an all-time high, with over 16 million people around the world clinically diagnosed with opioid use disorder (1). Of particular concern is opioid use among pregnant women, due in the most part to the rapid transfer of opioids across the placenta to the unborn child (2). At birth, the abrupt cessation of opioid delivery can lead to severe withdrawal, resulting in an array of neurological, physical and gastrointestinal symptoms, and even death (3). Improved antenatal diagnosis and postnatal care has advanced overall survival rates (4), however the longer-term implications of prenatal opioid exposure (POE) remain poorly understood, and no effective interventions are in place to prevent, mitigate or reverse the negative effects of POE.

Over a lifetime, the long-term effects of POE manifest as altered growth and developmental trajectories from infancy into adulthood. Deficits include delays in gross motor development (5) and cognitive functioning (5,6), as well as mental health issues later in life such as increased anxiety and impulsivity (7,8), development of addiction (8), and increased risk of suicidality (8). Neurologically, developmental delays have been linked to reduced brain size at birth (9) and reduced white matter connectivity (9,10). This may be mediated by immune dysregulation, evidenced by increased expression of inflammatory genes correlating with impaired myelination (11), decreases in peripheral immune markers (12) and increased risk of infection (13,14). More recently, POE has been linked to enteric and intestinal changes, such as transient reduced gut motility (15), and in adult opioid use disorder, microbiome dysregulation is a common symptom (16).

Interpretation of these findings, formulating direct links to POE and advancing therapeutic approaches is difficult due to the high co-morbidity of women with opioid use and poly drug use, pre-eclampsia, infection, and mental and physical ill-health (17,18). Furthermore, there is no standardized care protocol post-discharge (19), and POE is associated with an increased risk of child hospitalization for assault, maltreatment or overdose (20). To address these challenges, the rodent model of POE offers a valuable tool, recapitulating many of the core behavioral and neurological features of human POE, including cognitive impairment (21–24), and increased anxiety-like behavior (25,26). As in humans, juvenile POE rodents show evidence of less mature and organized white matter tracts (23), changes to innate and adaptive immune system function in the brain and periphery (22,23), and altered gut-microbiota composition (27,28). Therefore the rodent model offers a unique opportunity to study POE with a view to discovering novel therapeutic approaches.

To this end we have identified the short chain fatty acid (SCFA) sodium butyrate (NaB) as a potential treatment for POE, supported by several findings indicating that NaB may counteract the long-term symptoms of POE. First, chronic opioid exposure has been linked to decreased production of SCFA in both adult humans and rodent models of POE (16,27). Notably, decreased SCFA availability during development impairs myelination and immune function in rodent models of neonatal antibiotic exposure and germ-free animals (29,30). Second, in humans NaB intake is linked to cognitive function (31), and in rodents has been shown to improve memory following prenatal insult (30,32). Third, NaB reverses brain changes induced by perinatal insults, including rescue of de-myelination associated with neonatal antibiotic exposure (30), and improved white matter formation in a rat model of neonatal hypoxia-ischemia (33). And finally, NaB mediates enteric, peripheral and central nervous system immune function through regulation of cytokines and immune cell activation (34,35), and maintenance of gut- and blood brain-barrier integrity (36,37).

For these reasons, the aim of this study is to assess the efficacy of NaB to rescue impairments induced by prenatal exposure to methadone. We hypothesized that NaB treatment would mitigate the negative impacts of methadone on rat pups’ anxiety-like behavior, attention and cognition, brain development, peripheral inflammation, and that this would be mediated through altered gut-microbiome composition in pups and disrupted butyrate production in dams. To test this hypothesis, pregnant rat dams either received methadone via mini osmotic pump and NaB via drinking water in a 2 x 2 design. Methadone was chosen as it is the standard replacement therapy for women with opioid use disorder (38,39), with between 53 to 58% of pregnant women with OUD prescribed methadone or buprenorphine (40) to improve infant outcomes. We then performed a battery of developmental, behavioral, microbiome and neurobiological assessments across infancy, adolescence and adulthood that revealed NaB effectively rescued specific deficits associated with POE.

## Method

### Subjects

Twenty-six time-mated female Sprague Dawley rats (223-332g; Animal Resource Centre, AU) were purchased at gestational day 3 (GD3) and maintained on a 12-hour light cycle (on at 0700h) in a temperature- and humidity-controlled colony room (22°C±1) with *ad libitum* access to food.

All dams gave birth within 24 hours of GD22 (postnatal day 0) and litters were culled to 10 (5 male, 5 female where possible). On P22 pups were weaned into cages of 2-5 rats, matching for sex and experimental condition.

All procedures were approved by the University of New South Wales Animal Care and Ethics Committee and conducted in accordance with the NHMRC Australian Code for the Care and Use of Animals for Scientific Purposes (8^th^ Edition).

### Drugs

Methadone hydrochloride (National Institute of Measurement, NSW, AU) was dissolved in sterile saline and administered to dams via mini osmotic pumps (2ML2, Alzet CA, USA) at 9 mg/kg/day (41) from GD10 until P17 (Fig 1A). Minipumps were used to minimize the impact on the dams (42). This dose produces stable concentration in dam blood, accumulation in pup brain (43), and physiologically-relevant concentrations in pup urine (23).

**Figure 1. Dam microbiome composition and inferred butyrate production**

**A** Schematic of the experimental timeline indicating when fecal samples were collected from dams. **B** Microbial species richness by treatment group. **C** Shannon’s diversity by treatment group. Data as mean ± SEM with individual points overlaid. Data was analyzed using linear mixed effects models controlling for litter and cohort. *p<0.05 Tukey-corrected simple main effects examining pump by fluid interaction. **D** Principal-component analysis displays microbiome composition differences by treatment group in terms of beta-diversity (Aitchison distance; ellipses represent group 95% confidence intervals. **E** Targeted analysis of microbial genes involved in butyrate production. **F** Dam fecal butyrate concentrations. Data as mean ± interquartile range with individual points overlaid. Data was analyzed using linear mixed effects models controlling for litter and cohort. Percentages indicate observed distribution of 4 known pathways involved in microbial butyrate production. Sham/Veh n=6, Sham/NaB n=6, Met/Veh n=6, Met/NaB n=7. Asterisk, obelisk and hashtag indicate a significant main effect of pump, fluid or pump by fluid interaction respectively.

Sodium butyrate (Sigma-Aldrich, Missouri, USA) was administered via drinking water (3% w/v, refreshed every 48-72 hours) across GD10-P12. This concentration is well tolerated by rats and results in increased fecal levels of butyric acid (44). The period of administration was chosen to mimic treatment during gestation in human infants, as postnatal day 12 in rats is neurodevelopmentally equivalent to the end of trimester 3 in humans (45).

### Procedures

On GD10 dams were randomly allocated to groups: sham surgery/water (Sham/Veh), methadone pump/water (Met/Veh), sham surgery/sodium butyrate (Sham/NaB) and methadone pump/sodium butyrate (Met/NaB). Mini-osmotic pump surgery was performed according to manufacturer’s instructions (46) at GD10 and replaced at P3 to permit continuous methadone exposure across gestation and the early postnatal period (GD10 until P17; Fig 1A). Control animals received sham surgery as sham and saline-filled mini pumps insertion produce similar effects (41,47).

Maternal behavior was observed (P1-P10) at 2-minute intervals for 30 minutes/day in the dams’ home cages ((48); Supplementary Methods i). The percentage of pups in each litter with both eyes open recorded.

### Open-Field Test (P24)

To assess anxiety-like behavior one female and one male from each litter underwent the open-field test on P24. Each pup was placed in the center of a Perspex arena (L 60cm x W 60cm x H 50cm) for 5 minutes. Pose estimation was performed using Deeplabcut 2.3 (49); locomotion, time and entries in regions of interest, and rearing duration and frequency were scored using SimBA 1.5 (50).

### Adult Behavior and Cognition Testing (∼P65)

At ∼P65 one female and male pup from each litter were allocated for each behavioural test.

### Trial Unique Non-Matching to Location (TUNL) Task

The TUNL task of spatial and working memory was performed in standard trapezoid CANTAB boxes (Campden Instruments, UK) delivering sucrose pellets as reward (45MG Dustless Precision Sucrose Pellet, Able Scientific, AU). The protocol was consistent with past studies (51). Training, TUNL and delay tasks are described in full in Supplementary Methods ii.

### Five Choice Serial Reaction Time Task (5-CSRTT)

The 5-CSRTT procedures were based on Bari et. al 2008 (52) and tests motor impulsivity and sustained attention. Methods are described in full in Supplementary Methods iii, including training followed by variation of stimulus duration or inter-trial interval.

### Tissue Processing and Immunofluorescence (P21)

On P21 a naïve female and male pup from each litter were perfused with 4% paraformaldehyde, post-fixed for 24 hours, transferred to 70% ethanol and then paraffin-embedded (Biospecimen Preparation Laboratory, UNSW). Brains were sectioned (5µM) and immunofluorescence performed to detect microglia (rabbit polyclonal anti-iba1 antibody 1:100, 109-19741; Novachem, AU), astrocytes (mouse monoclonal anti-GFAP antibody 1:200; MAB360, Merck Millipore, GER), and myelin basic protein (1:500, M3821; Merck Millipore). Full details of the protocol are in Supplementary Methods iv.

Sections were imaged at 20x magnification using a slide scanning epifluorescence microscope (Axio Scan.Z1, Zeiss, GER) with ZenPro Software (Zeiss) and analyzed with ImageJ (National Institutes of Health, Bethesda, US), details of morphological analysis are in Supplementary Methods vii. Details of brain regions imaged are available in Supplementary Methods Fig. ii.

### Real-time Quantitative Polymerase Chain Reaction (RT-qPCR) (P21)

At P21 one naïve female and male pup from each litter were sacrificed and brains snap frozen in liquid nitrogen until processing. Using a brain matrix, punches were collected from 1 mm slices across the striatum (2 mm diameter) and dorsal hippocampus (1 mm diameter) into TRIzol (ThermoFisher Scientific, USA), see details of brain regions Supplementary Methods Fig. iii. Total RNA was isolated according to the manufacturer’s instructions. RNA was treated with DNAse1 (Merck, Australia) and reverse-transcribed using the iScript cDNA Synthesis Kit (Bio-Rad Laboratories, California, USA).

RT-qPCR gene expression analyses were performed in a StepOnePlus^TM^ System (Applied Biosystems, Massachusetts, USA) using gene-specific TaqMan assays, summarized in Supplementary Methods Table iii. All samples were run in triplicate and the expression of target genes normalized against housekeeper genes *Gapdh* and *Ubc*. Statistics were performed on expression levels and data represented as fold change relative to Sham/Veh.

### Cytokine Multiplex

At P7 and 21 blood was collected from the perfusion circuit and serum isolated to assess levels of 23 cytokines and chemokines using a magnetic bead immunoassay (Bio-Plex Pro™ Rat Cytokine 23-Plex Assay CAT #12005641, Bio-Rad Laboratories, California, USA), see Supplementary Methods v and Supplementary Methods Table iv for details. The plate was read using a Magpix Luminex machine, and median fluorescence data were collected. Cytokine concentrations were calculated using Milliplex Analyst software based on standard curves per cytokine.

### Microbiome Analysis

Fecal samples were collected from dams at GD10, P3 and P22, and from pups at P22, and the conclusion of adult behavioral testing (approx. P120). DNA was extracted using the QIAamp Fast DNA Stool Mini Kit and QIAamp DNA Mini Kit (QIAGEN, Hilden, Germany). Metagenomic library preparation with Illumina library prep, and sequenced on the NovaSeq X Plus platform (Illumina, San Diago, USA) at the Ramaciotti Center for Genomics (UNSW, Sydney, Australia). Sequencing adaptors and low-quality sequences (phred quality ³ >20) were trimmed using fastp v0.20.0 (53). PolyG tail trimming and base correction by overlap analysis was also performed using fastp with options --trim_poly_g and --correction, respectively. Tr Sequencing adaptors and low-quality sequences (phred quality ³ >20) were trimmed using fastp v0.20.0 (53).immed reads were mapped to the rat genome (GRCr8) using bowtie2 v2.5.2 (54) and the unmapped paired-end reads were extracted using samtools v1.19.2 (55). Cleaned non-host reads were used in taxonomic and function profiling. Taxonomic profiling of the bacteria was performed with MetaPhlAn4 following read alignment to the MetaPhlan4 database (56).

Functional profiling was conducted with HUMAnN 3.0 (57) using default parameters. QC’ed reads were mapped to UniRef90 gene families and MetaCyc metabolic pathways. UniRef90 gene-families were regrouped into KEGG orthology (KO) groups using the command ‘humann_regroup_table’. All functional annotations were normalized using the command “humann_renorm_table” to copies per million (--units cpm). To establish if the butyrate treatment was supplementing existing butyrate levels, or replacing diminished or disrupted butyrate production, we evaluated the distribution of genes involved in the four known butyrate pathways: Acetyl-CoA, 4-aminobutyrate/Succinate, glutarate and lysine.

SCFA quantification was performed with LC-MS on the QExactive (Thermo Scientific) Mass Spectrometer using the Agilent C18 1.9 m, 100 x 2.1 mm column at Monash Institute of Pharmaceutical Sciences and Monash Proteomics and Metabolomics Platform (MPMP) MIPS node (Monash University, Melbourne, Australia). Samples, spiked Quality control (QC) samples, and calibrants were extracted with acetonitrile and derivatized using 3-NPH. Isotopically labelled internal standards for SCFAs were derivatized using 13C6-3-NPH (58). Six SCFA derivatives were quantified: acetic (AA), propionic (PA), butyric (BA), valeric (VA), isobutyric (IBA) and isovaleric (IVA) acids (free acid form). Standard curve dilutions were prepared in 50% acetonitrile and spiked with internal standards. Final concentrations were calculated in ug/g dry weight.

### Fluorescein isothiocyanate-dextran (FITC) in vivo intestinal permeability test

A separate cohort of POE pups (Supplementary Methods vi) underwent assessment of intestinal permeability. On P9 pups were removed from their dam and housed on a heating pad with their litter mate for two hours prior to gavage. Following fasting pups were gavaged with 600mg/kg fluorescein isothiocyanate-dextran (FITC; 80mg/ml in sterile 0.9% saline) (catalog number FD4-1G, Sigma-Aldrich USA). After one hour blood was collected from the tail tip. Following 30 minutes at room temperature blood samples were spun at 4 degrees Celsius for 10 minutes, 1000xg, and serum collected. Serum was diluted 1:4 in PBS and FITC concentration in was analyzed in duplicates using a FLUOstar Omega microplate reader (BMG Labtech, Germany) with excitation λ 485 nm and emission λ 520 nm.

### Data Analysis

To avoid litter effects and account for potential non-independence between offspring (59) the unit of analysis for all statistical tests was the dam, with one male and one female pup from each litter randomly selected to be included in each analysis unless otherwise stated (see online data repository http://hdl.handle.net/1959.4/106186 for full details on pups assignment to experimental outcomes). Intra-litter correlation (ICC) was evaluated for a subset of outcomes by comparing models with and without litter as a random factor. Litter effects were non-significant across measures, indicating negligible within-litter clustering. Following this all physical, and behavioral data were analyzed using mixed model ANOVA (sex treated as a within-subjects factor). The specific n for each outcome and group is reported in the legend of each figure. Serum data was log-transformed before analysis. Scoring of the OFT and histological analysis were performed by a blinded observer, while maternal care scoring was not blinded due to detectable differences between groups (e.g. the presence of a minipump, odor of NaB). For microbiome analysis, alpha diversity was assessed by calculating richness and the Shannon index in the vegan r package. Microbiome count data was assessed following a centered log-ratio (CLR) transformation. Beta diversity was calculated using relative abundance data in terms of Aitchison distance to generate principal component analyses (PCA) plots. Permutational multivariate analysis of variance (PERMANOVA) tests were conducted using the *adonis* function (*vegan* package) to determine if beta diversity was significantly different between treatments. All microbiota measures were assessed using a linear mixed effects model approach (lme4 algorithms in tjazi package in R (60)), controlling for litter and cohort effects. Differential abundance analysis was corrected for multiple testing (false discovery rate (FDR) q < 0.05).

For exploratory analysis of the relationship between infant myelination and adult behavior, and maternal care and adult behavior, multiple correlations were correcting using the Benjamini-Hochberg FDR procedure. Significant effects were defined as FDR-adjusted p<0.05.

The study has ∼6% power to detect a small interaction (f = 0.10), ∼13% power to detect a medium interaction (f = 0.25), and ∼29% power to detect a large interaction (f = 0.40); the minimum interaction detectable with 80% power is f ≈ 0.74 (η^2^ ≈ 0.36). As such non-significant sex by treatment interactions do not rule out the presence small or moderate interaction effects.

Unless otherwise stated data were analyzed using IBM SPSS Statistics 26 software package. For all analyses rejection of the null hypothesis was set at α=0.05. Partial eta squared (η_p_^2^) values were reported as a measure of effect size, where a η_p_^2^ of 0.14 was considered a large effect size. Data were graphed using GraphPad Prism10. All significant statistical results for the supplementary data are contained in Supplementary Statistics.

## Results

Summary data on dam and pup weight gain across treatment, maternal care and eye opening is available in Supplementary Results Fig. iA-D and iv. NaB treated dams gained weight at a slower rate across gestation (Supplementary Results Fig. iB) and consumed more water across the postnatal period (Supplementary Results Fig. iC). Final NaB intake averaged 6.221g/kg/day, with the pattern of NaB consumption differing between Met/NaB and Sham/NaB across the gestational but not the postnatal period, and did not alter total NaB exposure (Supplementary Results Fig. iD). GI function was recorded during post-surgery monitoring from G10-17 and P3-10 and no disruption was observed.

### Methadone Exposure Dysregulates Microbial Functional Genes Required for Butyrate Production in Dams

Dam gut microbiota composition and function at the species-level were assessed by sequencing the fecal microbiome during exposure to methadone and NaB across gestation and lactation (Fig 1A). Microbial species richness was significantly reduced in both methadone-treated groups compared to Sham/Veh controls (Fig 1B; pump by fluid interaction; F_1,95_=5.429, p=0.022; post-hoc p<0.05). Shannon’s diversity (Fig 1C) did not differ. Microbiota beta diversity increased with methadone exposure and the extent of this depended on NaB treatment (Fig1D; pump by fluid; F_1,75_=1.371, p=0.008, R^2^=0.013, ω_p_^2^=0.004) by PERMANOVA after controlling for litter and cohort. Further assessment of individual species differences identified 40 species that were significantly altered by Methadone, NaB or both (Supplementary Results Fig. ii).

Since we hypothesized that methadone exposure would disrupt microbial butyrate production in the dam microbiome (61,62), we examined functional microbial genes directly involved in butyrate production (Fig 1E). All but 2 of the 17 genes involved in the four butyrate synthesis pathways were identified. Of the 15 genes, 6 were significantly altered by methadone, NaB or their interaction ((R,R)-butanediol dehydrogenase (K00004): drug effect: F_1,29.8_=9.078, p=0.003; 3-hydroxybutyrate dehydrogenase (K00019): liquid effect: F_1,29.8_=4.909, p=0.034; acetate CoA (K01034): drug by day: F_1,77.5_=2.912, p=0.027; glutaconate CoA-transferase, subunit B (K01040): drug effect: F_1,28.6_=6.392, p=0.019; 3-keto-5-aminohexanoate cleavage enzyme (K18013): drug by day: F_1,81.3_=2.900, p=0.027; K19709: drug by day: F_1,24.5_=7.055, p=0.009).

As most identified genes (63.9%) were terminal genes indistinguishable between butyrate synthesis pathways, MetaCyc was used to map to quantify the distribution of butyrate synthesis potential between pathways. A total of 6 intermediate pathways were identified: 2 from the Acetyl-CoA (66.5%), 3 from the 4-aminobutyrate/Succinate pathway (32.8%) and 1 from the lysine pathway (0.74%) (Fig. 1E). To investigate whether these genomic changes were associated with changes in absolute butyrate expression, we examined fecal levels of SCFA. Butyrate was significantly decreased following methadone exposure (Fig 1F; F_1,82_=6.368, p=0.013), as were isobutyrate and valerate. In contrast, acetate was increased with methadone, and propionate was reduced by NaB exposure (Supplementary Results Fig. iii).

### Methadone Exposure Transiently Dysregulates Functional Microbial Genes Required for Butyrate Production in Pups and Increases Gut Permeability and is Reversed by NaB

We assessed pup microbiota composition and predicted altered butyrate production both at weaning and following adult behavioral testing (Fig 2A). Both microbial species richness (Fig 2B; pump by fluid by day by sex: F_1,84.7_=7.263, p=0.008) and Shannon’s diversity (Fig 2C; pump by fluid by day: F_1,101.3_=10.869, p=0.001; pump by fluid by sex: F_1,85.3_=6.408, p=0.013) were significantly altered by treatment, developmental day and pup sex. Notably, Shannon’s diversity was increased in Sham/Veh offspring across development (Tukey-adjusted posthocs; p=0.021 and p=0.012 respectively) with no differences over time detected in the other treatment groups. At PND22, Sham/Veh and Met/Veh pups were significantly different (p=0.014) - offspring exhibited a significant interaction between treatments and developmental day in microbiota composition (Fig 2D; F_1,100_=2.018, p=0.021, R^2^=0.015, ω_p_^2^=0.009), although this may be due to differences in dispersion (F =2.423, p=0.024). We assessed individual species differences and found 79 were significantly altered by the interaction between development day and methadone, NaB or both treatments (Supplementary Results Fig. v).

**Figure 2.**
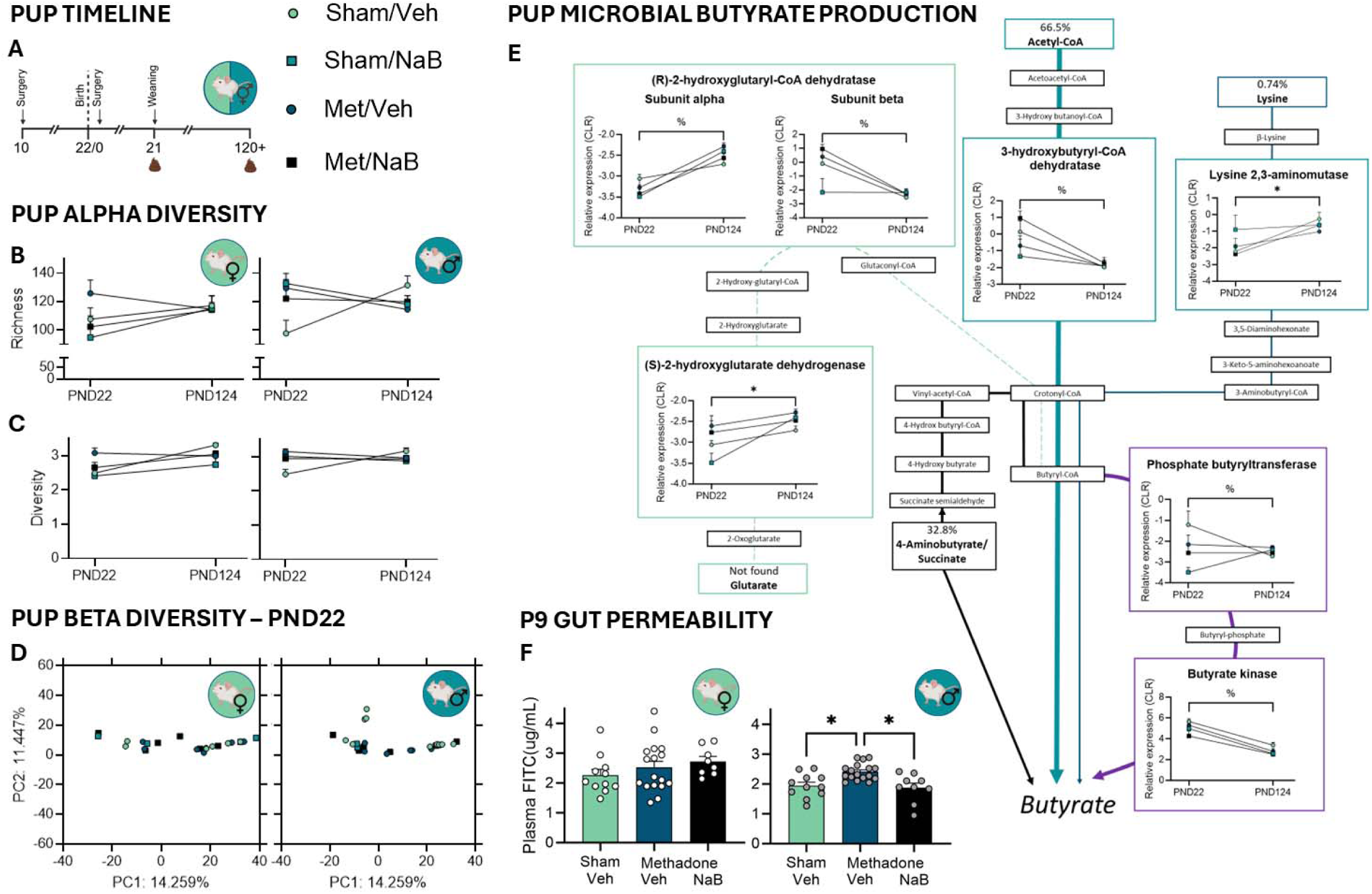
Offspring microbiome composition and inferred butyrate production. **A** Schematic of the experimental timeline indicating when fecal samples were collected from dams. **B** Microbial species richness by treatment group. **C** Shannon’s diversity by treatment group. Data as mean ± SEM with individual points overlaid. Data was analyzed using linear mixed effects models controlling for litter and cohort. *p<0.05 Tukey-corrected simple main effects examining pump by fluid interaction. **D** Principal-component analysis displays microbiome composition differences by treatment group in terms of beta-diversity (Aitchison distance; ellipses represent group 95% confidence intervals. **E** Targeted analysis of microbial genes involved in butyrate production. **F** In-vivo assessment of gut permeability at P9. Data as mean ± interquartile range with individual points overlaid. Data was analyzed using linear mixed effects models controlling for litter and cohort. P21 females Sham/Veh n=6, Sham/NaB n=6, Met/Veh n=6, Met/NaB n=7 and males Sham/Veh n=6, Sham/NaB n=6, Met/Veh n=6, Met/NaB n=7. At P124 there were 2-4 pups/sex/litter included in analysis from Sham/Veh n=6, Sham/NaB n=6, Met/Veh n=6, Met/NaB n=7 litters, with litter controlled for using a linear mixed effects model. Specific numbers for each group are as follows – females Sham/Veh n=10, Sham/NaB n=5, Met/Veh n=8, Met/NaB n=9, males, Sham/Veh n=9, Sham/NaB n=5, Met/Veh n=8, Met/NaB n=12. Asterisk and percentage indicate a significant main effect of pump, or methadone, sodium butyrate and developmental day interaction respectively.

Furthermore, we examined functional microbial genes directly involved in butyrate production (Fig 2E). Of the 15 genes we identified, 7 were significantly altered by Methadone, NaB or their interaction (Phosphate butyryl transferase (K00634): drug by liquid by day effect: F_1,93.5_=10.026, p=0.002; butyrate kinase (K00929): drug by liquid by day effect: F_1,105.1_=9.003, p=0.003; beta-lysine 5,6-aminomutase alpha subunit (K01844): drug effect: F_1,113.5_ =4.313, p=0.040; (S)-2-hydroxyglutarate dehydrogenase (K15736): drug effect: F_1,112.3_=9.708, p=0.006; 3-hydroxybutyryl-CoA dehydratase (K17865): drug by liquid: F_1, 114.2_=7.440, p=0.007; (R)-2-hydroxyglutaryl-CoA dehydratase subunit alpha (K20903): drug by liquid by day effect: F_1,114.9_ =5.052, p=0.027; (R)-2-hydroxyglutaryl-CoA dehydratase subunit beta (K20904): drug by liquid effect: F_1,113.8_ =5.224, p=0.040). Interestingly, while microbial butyrate production in both dams and pups were impacted by our interventions, in pups these disruptions appear to impact specific pathways, rather than universal, terminal enzymes as observed in the dam microbiota.

To assess the functional impact of these changes in microbial genes we ran an *in vivo* assessment of gut permeability using FITC. On P9 male pups exposed to methadone alone demonstrated increased gut permeability compared to controls, and this was reversed in pups whose dam had been treated with NaB (Fig. 2F; F_1,34_=8.292, p=0.001, η_p_^2^=0.328; Tukey-adjusted post-hoc; p=0.008 and p=0.004 respectively).

### Methadone Increases Anxiety-like Behavior in Adolescence and this is Partially Mitigated by NaB

On P24 pups underwent the OFT (Fig. 3A). All methadone-exposed pups made fewer entries into the center of the field (Fig. 3C, F_1,19_=7.420, p=0.013, η_p_^2^=0.271) and methadone exposed females spent less time in the center area (Fig. 3D, F_1,21_=7.516, p=0.012, η_p_^2^=0.264), indicative of increased anxiety-like behavior. During the OFT, pups that have been exposed to methadone spent less time rearing which was rescued with NaB exposure (Fig. 3B, interaction effect: F_1,18_=5.000, p=0.033, η_p_^2^=0.218); rearing is a stress-sensitive measure (63) that is critical for spatial learning in young rodents (64). No effects on locomotion were observed. In summary, NaB partially rescued anxiety-like behavior, and may promote exploration and learning in a novel environment.

**Figure 3.**
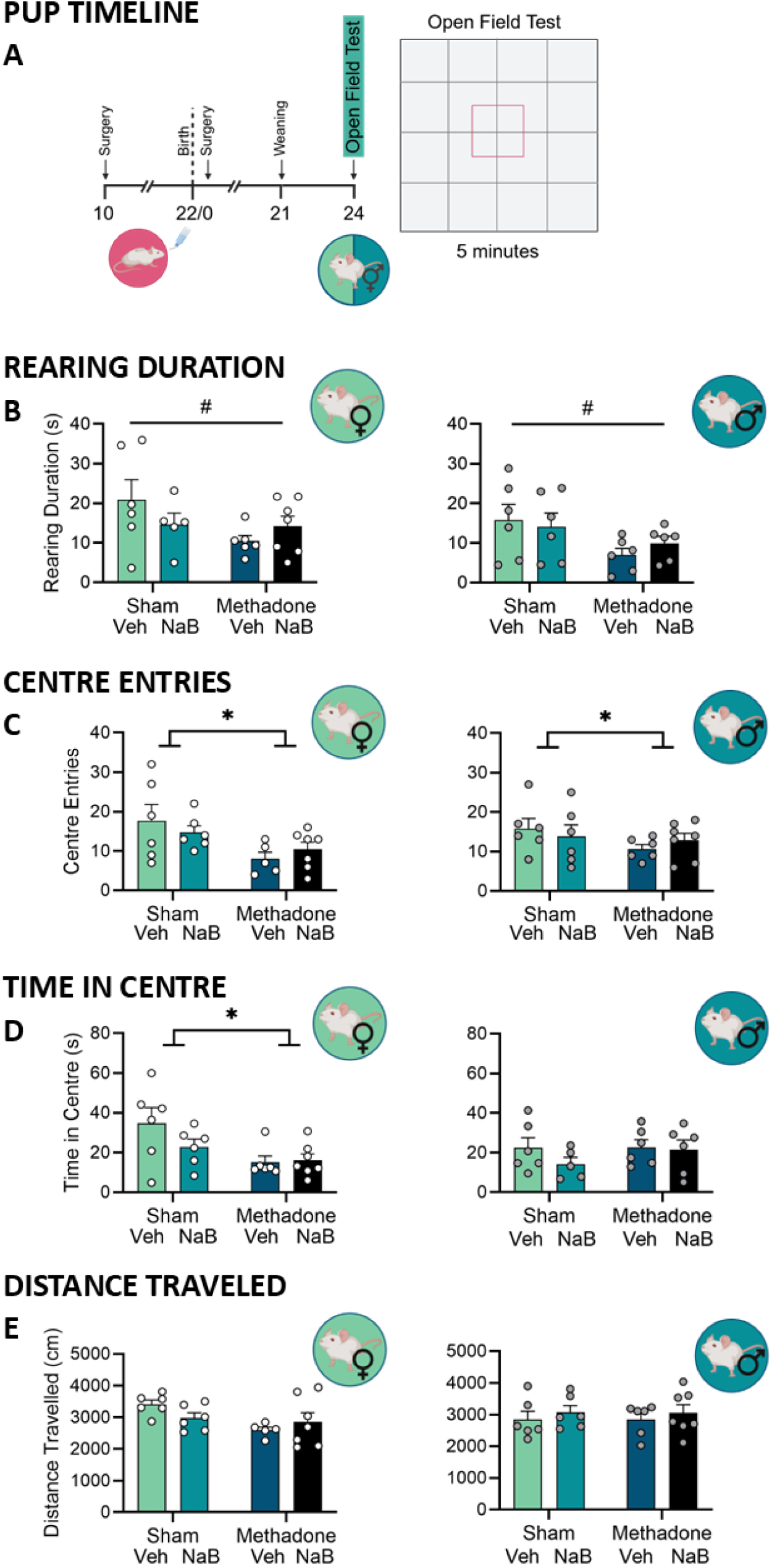
Pups performance during the open field test. **A** Schematic of the open field test which was performed on postnatal day 24. **B** The amount of time pups spent rearing, **C** the number of entries in to the center of the open field, **D** amount of time spent in the center of the open field and **E** distance travelled were all recorded using Deeplabcut 2.3 and SimBA 1.5. Data points represent group mean **±** SEM with female (white points, Sham/Veh n=6, Sham/NaB n=6, Met/Veh n=6, Met/NaB n=7) and male (grey points, Sham/Veh n=6, Sham/NaB n=6, Met/Veh n=6, Met/NaB n=7) pups. Asterisk and hashtag indicate a significant main effect of pump or pump by fluid interaction respectively.

### NaB Rescues Methadone-induced Deficits in Spatial Learning and Working Memory

A female and male pup from each litter underwent the TUNL task in adulthood to assess working memory and pattern separation (Fig. 4A). Female rats who had been exposed to methadone took longer to learn the TUNL task than females exposed to methadone and NaB (Fig. 4B, interaction effect: F_1,19_=5.307, p=0.034, η_p_^2^=0.238). On the first day of the TUNL task female rats exposed to methadone or NaB made fewer correct responses than female rats who had been exposed to methadone and NaB together (Supplementary Results Fig. viB). All groups took a similar number of days to reach delay testing criteria and made a similar number of correct responses in the final two sessions prior to delays being introduced (Supplementary Results Fig. vC). However, in these final sessions methadone-exposed animals touched the screen more during inter-trial intervals, indicative of increased impulsivity and habitual behavior (65), which was rescued in animals who were treated with NaB (Fig. 3C, interaction effect: F_1,19_=4.880, p=0.040, η_p_^2^=0.204).

**Figure 4.**
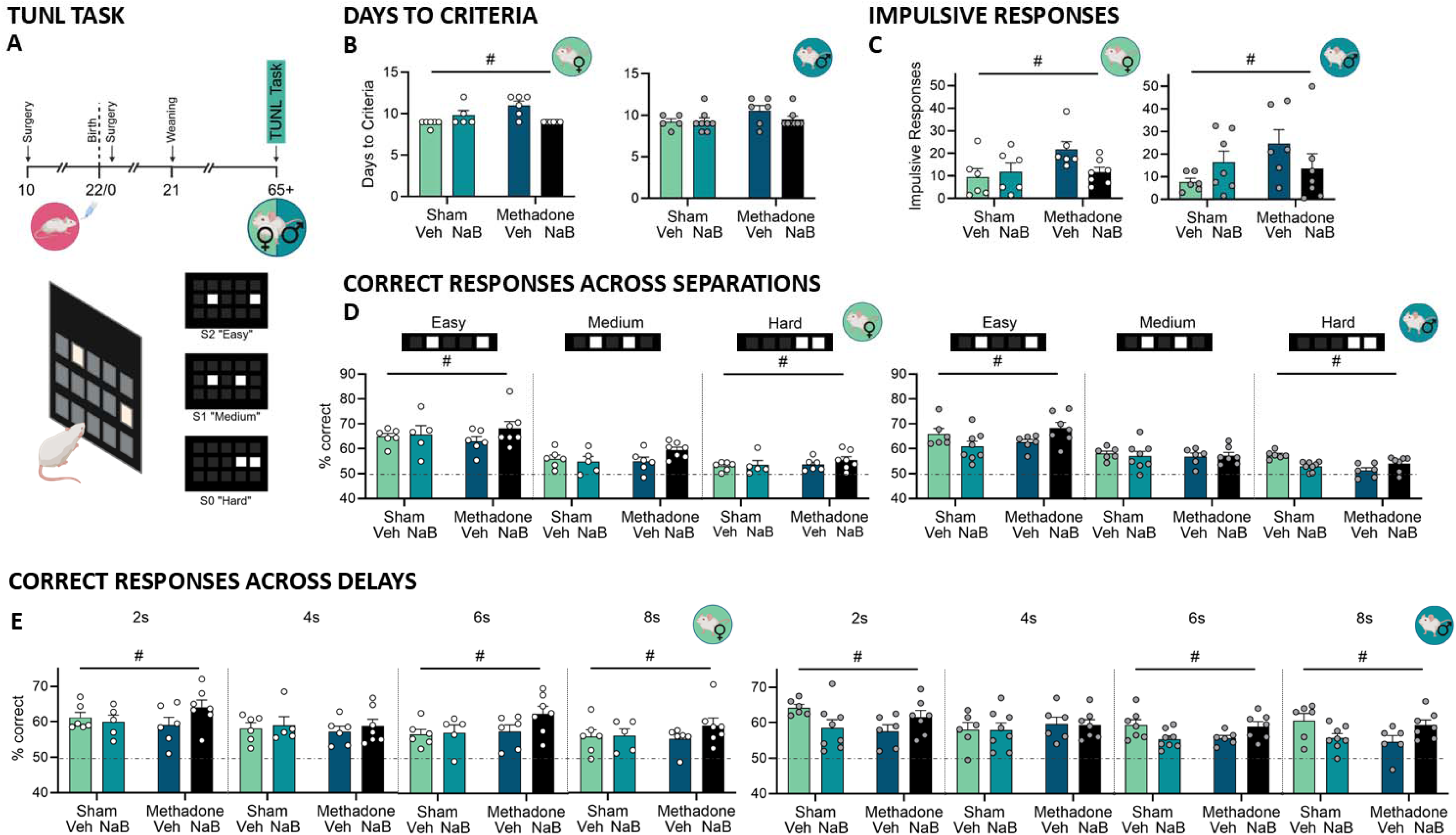
Performance on the trial unique non-matching location task. **A** Schematic of the TUNL task which one female and one male pup from each litter underwent from P65+. **B** The number of days taken for each rat to reach pretraining criteria and progress to the TUNL task. **C** The number of touches to the screen made during the inter-trial interval in the final TUNL training session before delays were introduced. **D** The percentage of correct responses made at “easy” (S2), “medium” (S1), and “hard” (S0) spatial difficulty trials across all delay sessions. **E** The percentage of correct responses made at each delay, 2 seconds, 4 seconds, 6 seconds or 8 seconds, across all spatial separations. Data points represent group mean **±** SEM with 6/7 female (white points Sham/Veh n=6, Sham/NaB n = 5, Met/Veh n=6, Met/NaB n=7) and male (grey points Sham/Veh n=6, Sham/NaB n = 8, Met/Veh n=6, Met/NaB n=7) pups/group. The dashed horizontal line represents performance at chance, with no preference for the novel location. Hashtags indicates a significant interaction effect.

Across delay sessions, rats exposed to methadone made fewer correct responses than those exposed to methadone and NaB together at both the “easy” and “hard” spatial difficulty level rats (Fig 4D interaction effect; “easy”: F _1,19_=8.331, p=0.009, η_p_^2^=0.305, “hard”: F _1,19_=6.086, p=0.023, η_p_^2^=0.243). When the delay was 2-, 6-, or 8-seconds, methadone or NaB exposed rats made fewer correct responses than rats that had been exposed to both methadone and NaB together (Fig. 4E, interaction effect; 2s delay: F_1,19_=4.797, p=0.041, η_p_^2^=0.202, 6s delay: F_1,19_=9.223, p=0.007, η_p_^2^=0.327, 8s delay: F_1,19_=5.284, p=0.033, η_p_^2^=0.218). An interaction between separation and delay across all sessions (F_6,114_=2.519, p=0.025, η_p_^2^=0.117) indicated that delay and spatial difficulty interacted to impact performance (see Supplementary Results Fig. vii).

In summary, perinatal methadone impaired working memory and pattern separation, and NaB treatment rescued these effects in both male and female rats. The effects of methadone were not dependent on pattern difficulty or task delay, indicating that methadone likely impacts universal behavioral processes underlying the task.

### NaB Rescues Methadone-induced Deficits in Sustained Attention and Impulsivity

A naïve male and female pup from each litter underwent the 5-CSRTT in adulthood to assess motor impulsivity and sustained attention (Fig. 5A). All rats reached criteria for multiple stimulus duration (MSD) and multiple inter-trial interval (MITI) sessions at a similar rate (Supplementary Results Fig. viii). Performance as indicated by perseverative responses, response latency and reward collection latency, was similar across all groups in the final training session (Supplementary Results Fig. ix) indicating a similar level of motivation to obtain the sucrose reward.

**Figure 5.**
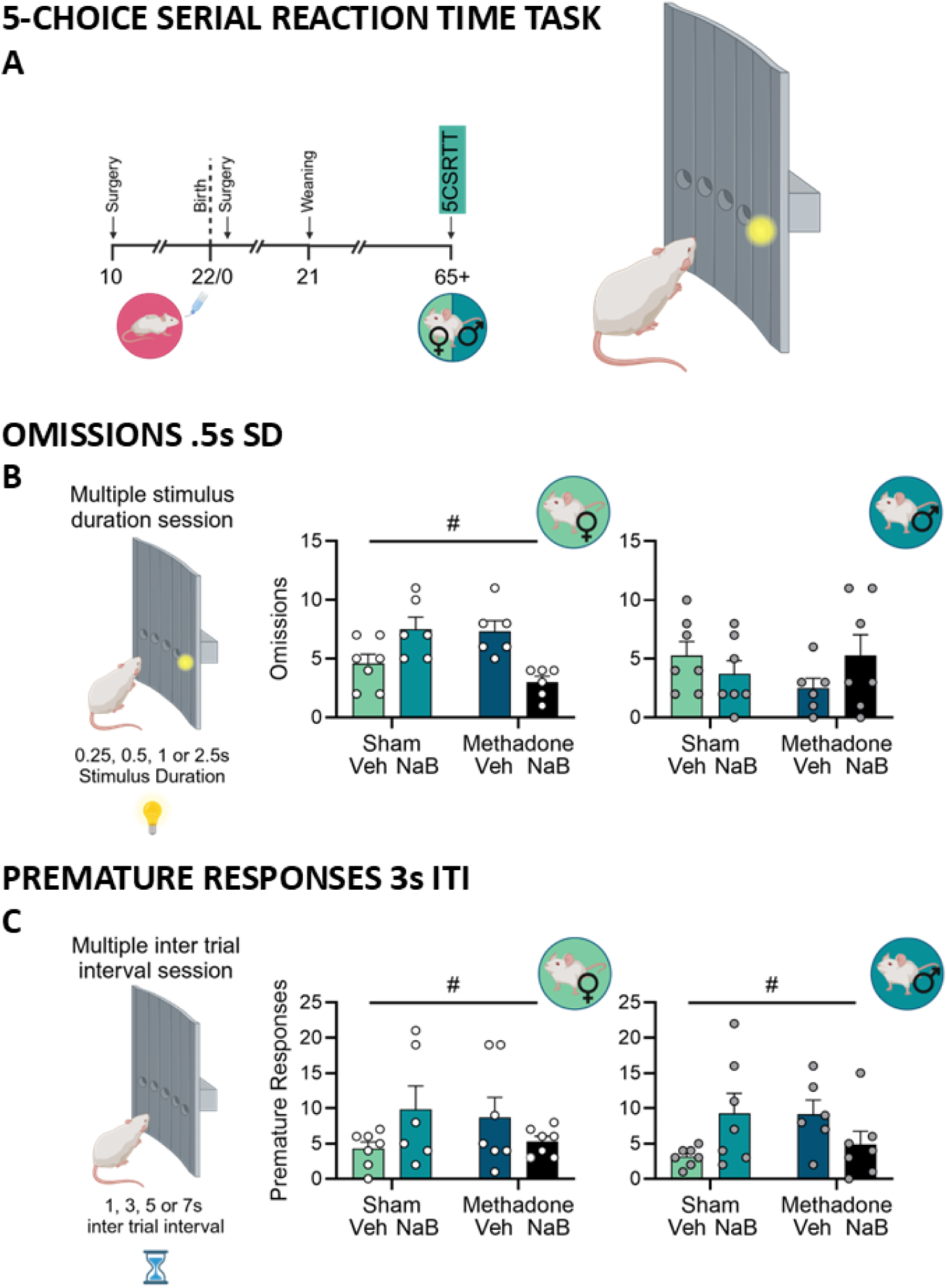
Performance on the 5-choice serial reaction time task. **A** Schematic of the 5-choice serial reaction time task, which one female and one male pup from each litter underwent from P65+. **B** The number of trials where rats failed to respond (omissions) when the stimulus durations (SD) was .5 seconds in the multiple stimulus duration (MSD) session. **C** The number of responses inappropriately made before the stimulus was displayed (premature responses) when the intertrial interval (ITI) was 3 seconds during the multiple inter trial interval (MITI) session. Data points represent group mean **±** SEM with 6/7 female (white points Sham/Veh n=7, Sham/NaB n=6, Met/Veh n=6, Met/NaB n=7) and male (grey points Sham/Veh n=7, Sham/NaB n=7, Met/Veh n=6, Met/NaB n=7) pups/group. Hashtags indicate a significant interaction effect.

Once rats had reached training criteria, they underwent the MSD session, with pseudo randomly varied SD of 0.25, 0.5, 1, 1.25 or 2.5 seconds, where a shorter SD requires higher levels of sustained attention. When the SD was at its second shortest durations, 0.5s, female rats who had been exposed to methadone missed more trials than if they had additionally been exposed to NaB (Fig. 5B interaction effect: F_1,20_ =15.556, p<.001, η_p_^2^=0.414), suggesting impaired sustained attention following methadone that is rescued with additional NaB exposure. Performance across other SDs was similar between groups. Overall, male methadone exposed rats were quicker to make incorrect responses than sham males (Supplementary Results Fig. xD), while perseverative responses, correct and incorrect responses, and reward collection latency were similar across groups (Supplementary Results Fig. x).

Rats progressed to the MITI sessions where the ITI varied between 1, 3, 5 and 7 seconds, with a longer ITI requiring rats to withhold a disadvantageous impulsive response for a longer period of time. When the ITI was 3s rats that had been exposed to methadone made more premature responses than those who had been exposed to both methadone and NaB (Fig 5C, interaction effect: F_1,20_=5.904, p=.024, η_p_^2^=0.212) suggesting that methadone exposure increased impulsive action, which was rescued by exposure to NaB. Performance across other ITIs was similar, and perseverative responses, correct and incorrect responses, and reward collection latency were similar across groups (Supplementary Results Fig. xi). Together these results suggest subtle and sex-dependent negative effects of methadone on sustained attention and motor impulsivity that can be rescued with NaB treatment.

### NaB Rescues Methadone-induced Decreases in Myelin Basic Protein Staining and Differences in Oligodendrocyte Expression

On P21 one female and male rat from each litter were perfused (Fig. 6A). Their brains were paraffin embedded and stained for MBP in the striatum and dHPC (Supplementary Methods Fig. iii) to give an indication of myelination in regions relevant to working and spatial memory (66), and motor impulsivity (67).

**Figure 6.**
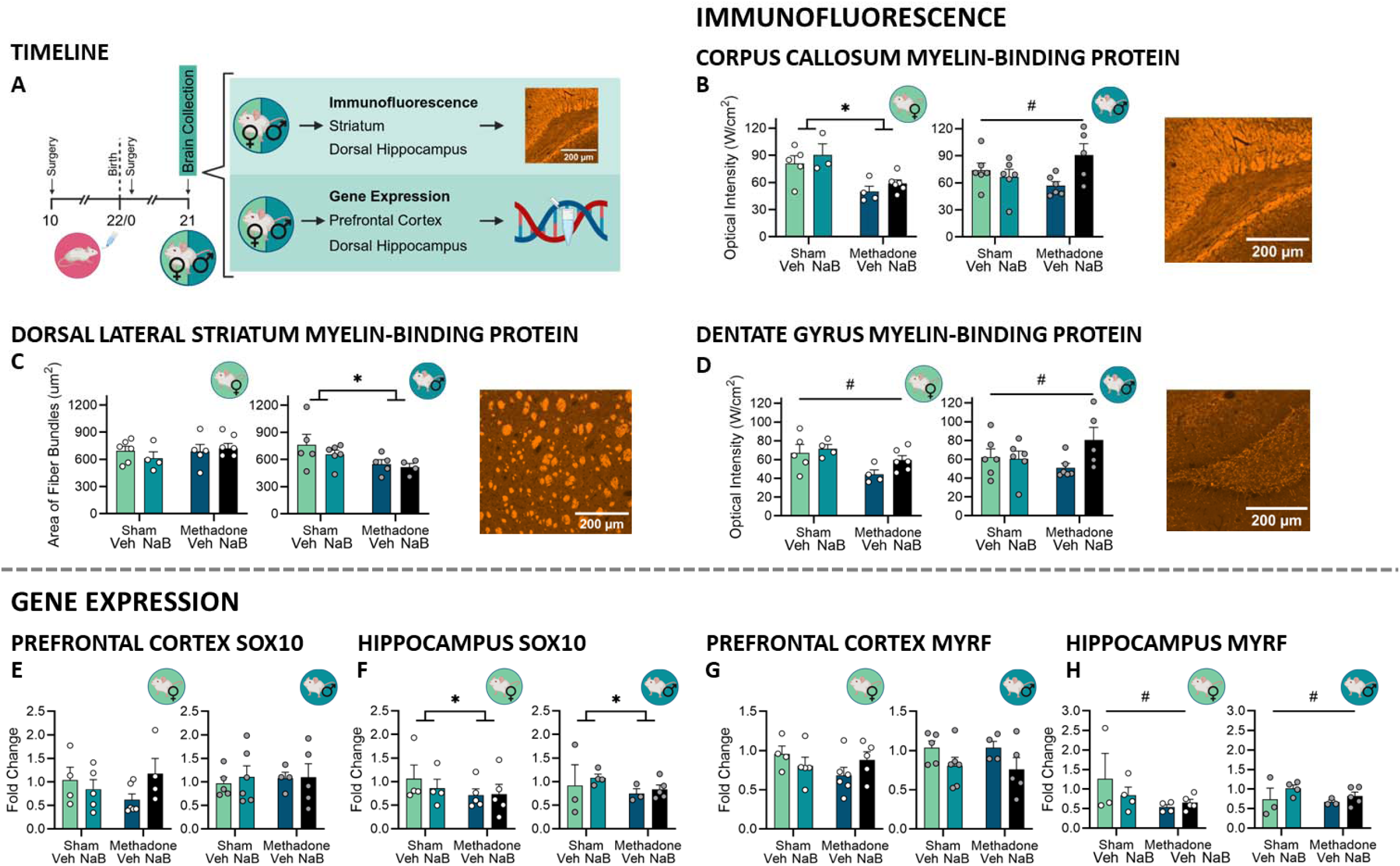
Analysis of myelination on P21. **A** Timeline of brain collection for immunofluorescence and RT-qPCR. **B** Intensity of staining for myelin-basic protein (MBP) in the corpus callosum at the level of the dorsal hippocampus and a representative image of MBP staining in this region. **C** Intensity of staining for MBP in the dorsal lateral striatum and a representative image of MBP staining in this region. **D** Intensity of staining for MBP in the dorsal hippocampus and a representative image of MBP staining in this region. Fold change expression of **E** *Sox10* in the prefrontal cortex, and **F** hippocampus, and of **G** *Myrf* in the prefrontal cortex, and **H** hippocampus. Data points represent group mean **±** SEM with 3-6 female (white points, immunofluorescence in the striatum Sham/Veh n=5, Sham/NaB n = 4, Met/Veh n=5, Met/NaB n=7 and dHPC Sham/Veh n=5, Sham/NaB n=4, Met/Veh n=4, Met/NaB n=6, and RT-qPCR in the PFC Sham/Veh n=5, Sham/NaB n=5, Met/Veh n=6, Met/NaB n=5, and dHPC Sham/Veh n=4, Sham/NaB n=4, Met/Veh n=5, Met/NaB n=5) and male (grey points immunofluorescence in the striatum Sham/Veh n=4, Sham/NaB n = 6, Met/Veh n=5, Met/NaB n=4 and dHPC Sham/Veh n=6, Sham/NaB n=6, Met/Veh n=6, Met/NaB n=5, and RT-qPCR in the PFC Sham/Veh n=5, Sham/NaB n=6, Met/Veh n=4, Met/NaB n=5, and dHPC Sham/Veh n=3, Sham/NaB n=5, Met/Veh n=3, Met/NaB n=5) pups/group. Asterisks and hashtags indicate a significant main effect of methadone or interaction effect respectively.

In the dentate gyrus, methadone and NaB exposed pups had greater staining intensity than those exposed to methadone alone (Fig. 6C interaction effect: F_1,14_=7.955, p=.014, η_p_^2^=0.4362). A similar pattern was evident in the corpus callosum, where male pups exposed to methadone and NaB had greater MBP staining intensity than those exposed to methadone alone (Fig. 6B interaction effect: F_1,19_=6.002, p=.024, η_p_^2^=0.240), while all females exposed to methadone had decreased staining intensity (F_1,14_=17.335, p=.001, η_p_^2^=0.553). In the dorsal lateral striatum male pups exposed to methadone had smaller fiber bundles (Fig. 6D: F_1,16_=5.815, p=.028, η_p_^2^=0.297). Overall these findings suggest that methadone exposure disrupts myelination in regions critical for working and spatial memory, and motor impulsivity. Co-administration of NaB can partially rescue these effects in a sex-specific manner.

An additional female and male pup from each litter were euthanized and brains collected for gene expression analysis by RT qPCR to assess changes in oligodendrocyte proliferation that may have contributed to alterations in myelination (Fig 6A). In the PFC female pups that had been exposed to methadone and NaB had increased *Sox10* expression compared to pups who had been exposed to methadone alone (Fig. 6E; sex by pump by fluid - F_1,13_=6.252, p=.027, η_p_^2^=0.325). In the hippocampus methadone exposed pups had decreased *Sox10* expression (Fig. 5F; F_1,6_=8.379, p=.032, η_p_^2^=0.563). In the hippocampus pups who had been exposed to methadone and NaB had increased *Myrf* expression compared to pups that had been exposed to methadone alone (Fig. 6G; F_1,6_=20.765, p=.004, η_p_^2^=0.776). However, there were no differences in *Myrf* expression in the PFC (Fig. 5H). MBP expression and tissue from the striatum were also analyzed and showed no differences in expression between groups (Supplementary Results Fig.xiv-xvi). These results indicate region- and sex-specific effects of methadone and NaB exposure on oligodendrocyte related gene expression, suggesting methadone may disrupt myelination by altering oligodendrocyte proliferation, an effect partially recused when NaB has also been administered.

### NaB, but not Methadone, Impacts Central and Peripheral Immune Activity

To assess potential impacts of methadone on immune function, central and peripheral inflammation was assessed using immunofluorescence staining for Iba1 and GFAP in the striatum and dHPC at P21, RT-qPCR assessing inflammatory gene expression in the PFC, striatum and dHPC at P21, and a cytokine multiplex assessment of blood serum on P7 and P21. These outcomes were minimally impacted by methadone exposure. However, NaB exposure increased the number of Iba1-positive cells in the corpus callosum, elevated CD68 and IL10 expression in the prefrontal cortex, reduced CD68 expression in the striatum, and altered cytokine serum levels. Together these finding suggests NaB mediated central and peripheral immune activity. These results are summarized in Supplementary Results Figures xii-xvi, and Supplementary Results Table i and ii.

## Discussion

Here we show that methadone exposure during pregnancy and lactation disrupts both maternal and offspring microbiota composition and microbial butyrate pathways in a rat model of POE, reduces dam fecal butyrate and increases gut permeability in male offspring. Furthermore, methadone exposure is associated with anxiety-like behavior and disrupted myelination during adolescence, and cognitive deficits in pattern separation, working memory and attentional processing in adulthood. We show for the first time that oral NaB treatment rescued methadone-induced deficits in spatial and working memory and reduces motor impulsivity. Molecular analysis suggests that the beneficial effects of NaB supplementation may be due to replacement of disrupted butyric acid supply in the maternal microbiota which supports offspring myelination. These data indicate that early life NaB has potential as a therapy to mitigate POE-induced deficits likely to impair academic performance and quality-of-life.

Poor cognition, a primary adverse outcome of POE, is itself associated with negative life outcomes including poor mental health, and increased risk of addiction and suicide in adulthood (8,68). Here we show for the first time that a simple, cheap and accessible dietary supplement administered across the perinatal period can significantly improve cognition in adulthood. Sodium butyrate reversed methadone-induced deficits in the TUNL task indicative of poor cognitive flexibility, working memory and spatial separation as well as impulsive action in both the TUNL and 5-CSRTT, to the extent that MET/NaB rats were indistinguishable from controls. These methadone-induced deficits are consistent with disruption across hippocampal and prefrontal circuitry, similar to that reported following POE (22,41) and in adults with opioid use (69–71). The finding that this simple treatment improves adult outcomes in these animals is a significant first step towards treating POE in humans, where no interventions currently exist. It is important to note here that some behavioral changes in the Veh/NaB group were detected, consistent with findings that disruption of gut health or gene expression in healthy individuals is not always beneficial (72,73), an important consideration for future studies suggesting NaB may not be universally beneficial in the absence of POE.

When considering possible mechanisms through which the rescue effect on behavior occurs, our results highlight several interacting possibilities. Endogenous opioid signaling plays a key role in stem cell differentiation and proliferation early in brain development (71), in particular oligodendrocyte differentiation (75). Previous human and animal research has highlighted poor white matter integrity as a consistent feature of POE (9,10,23,41,76). Similarly, butyrate has shown benefit for the prevention of demyelination or promotion of remyelination following neonatal insult (30,77). Here we hypothesize that cognitive and behavioral improvements observed with NaB in adulthood may be mediated, at least in part, through normalization or support of myelin formation in adolescence – a peak period of myelination (78). In adolescence and in a drug-free state, we found NaB rescued MBP staining intensity in the dentate gyrus of the hippocampus, a region critical for performance of the TUNL task (79). Our gene expression data further suggest this may be due to lasting opioid-induced disruption of oligodendrocyte differentiation (76) as evidenced by downregulation of *Sox10* following methadone exposure in adolescence, and a reversal of methadone-induced *Myrf* suppression with NaB co-treatment. As *Sox10* regulates *Myrf*, and both are essential for successful oligodendrocyte maturation and myelination (80), disruption of these pathways may lead to impaired myelination and connectivity in brain regions critical for cognition. Positive correlations between MBP intensity in adolescence and adult behavioral performance (Supplementary Results Fig. xxiii) supports the link between myelination and long-term outcomes, primarily in females. Future studies will explore NaB treatment during adolescence, a critical period for myelination (78), and a useful window for intervention, as well as more in-depth analysis using electron microscopy and diffusion tensor imaging.

A second interacting possibility is the role of gut health. Chronic opioid use has been associated with disrupted gut microbiota composition and reduced short-chain fatty acid production in adult users, likely due to reduced gut motility (16), and with a consequence of reducing gut barrier integrity (16), likely triggering both systemic (16) and neuro-inflammation (81). We show for the first time that microbial functional genes required for butyrate production are dysregulated by maternal opioid exposure in both the maternal and early offspring microbiota, and results in decreased dam fecal butyrate content. Additionally, we observed changes in a number of bacteria linked to inflammation and neurological issues (82,83). We hypothesize that replacement of NaB through oral administration counteracts this processes by enhancing gut barrier integrity (30,37), possibly due to NaB increasing expression of tight junction proteins (84). Indeed, male POE pups showed increase gut permeability that was rescued when pups had additionally been treated with NaB. This highlights a link between altered gut microbiota composition, metabolite production and gut permeability following POE, and the potential of NaB supplementation.

Although here we did not see changes in immune system function as result of methadone, NaB alone impacted on circulating cytokines, microglia count and the expression of immune cell genes was detected. This pattern is consistent with previous reports showing NaB enhances innate immune signaling and cytokine production (85,86), and upregulates anti-inflammatory mediators such as IL-10 (87). The possibility of NaB altering microglial state and thus facilitating neural development, was further explored using morphological analysis of microglia in the dentate gyrus (Supplementary Methods vii), although no group differences were detected (Supplementary Results Table iii). An interaction of methadone and NaB was not evident at this single early time point, however it is possible that this did not align with the temporal pattern of methadone-induced changes, with further investigation warranted given the dynamic nature of microglial state (88).

A key feature of NaB is its well defined role as a histone deacetylase inhibitor (iHDAC) (77). Through this action NaB is thought to support myelination (77), enhance gut (89) and blood brain barrier function (90), and modulate immune function (91). As POE induces epigenetic modifications to chromatin (92), it is likely that epigenetic regulation of gene expression across multiple organ systems plays a role in these outcomes, and in particular enables persistent changes in plasticity-related genes critical to learning and memory that extend well beyond the last drug treatment. Exploring this possibility and how it may interact with brain development and gut health is the focus of ongoing studies in our laboratory.

Future experiments will alter NaB administration pathway, reducing exposure of the gut to NaB, whilst also tracking markers of gut microbiota composition and gut permeability. Improved neural development and behavior in the absence of gut alterations would support epigenetic regulation as a primary pathway for NaB mediated improvement. Alternatively, probiotics known to improve gut barrier function and reduce bacterial translocation without altering gene expression, such as *lactobacillus* (93), could be administered. If improvements in gut barrier function and behavior are observed free from the need for epigenetic regulation this would suggest the positive impact of NaB is due to its post-biotic properties. Further experiments will help to identify the mechanism of NaB’s primary therapeutic impact and allow the development of more targeted treatments. It is also likely that NaB’s ability to act upon multiple pathways in tandem is part of what makes its particularly appropriate for the treatment of opioid exposure, a drug class which also acts across many systems simultaneously.

When considering the translational potential of these findings, it is important to consider methodological factors that may influence outcomes. Here we use mini-osmotic pumps with sham controls, largely due to welfare issues associated with repeat injections that impact on our key dependent variables (94,95), yet acknowledging that the metabolic profile of rats and accumulation in the fetus does differ from humans (43,96). Further, we did not detect evidence of withdrawal in our methadone exposed pups (e.g. abrupt weight changes, behavioural change). This may be due to the continued delivery of methadone via lactation paired with the gradual decrease in exposure through to weaning, and is consistent with past studies finding no evidence of withdrawal-induced ultrasonic vocalizations when using a similar procedure (43). Another factor potentially impacted by drug delivery mode is maternal care. Here, using a simple method of maternal care assessment we detected reduced arch-backed nursing and licking and grooming in the methadone and NaB group which in some circumstances has been linked to altered behavioral outcomes (48). Although we cannot rule this out as a factor, there was no correlation between maternal care and key adult behavior outcomes (Supplementary Results Fig. xvii and xviii).

When considering capacity for translation, it is important to identify that the concentrations of both methadone and NaB used here fall within the dose range that is pharmacologically relevant to humans. Previous research indicates that the dose of methadone administered via minipumps produces stable plasma and urine concentrations comparable to that observed in pregnant women on opioid maintenance therapy (43). This extends into delivery via lactation, where the same dose of methadone via minipump across gestation and lactation results in methadone detected in the pups urine at P14 (23). For NaB, oral administration in rodents leads to rapid metabolism into acetyl-CoA, which then enters the Tricarboxylic acid (TCA) cycle (97). It has an effective half-life of less than five minutes (98). Despite this short half-life, treatment with NaB across gestation and lactation in mice has been shown to lead to increase serum NaB levels over 12 days after exposure ended (99). In humans, the pharmacokinetic profile is similar, with a half-life of 15-30 minutes in circulation (100), necessitating frequent administration. Although species differences in diet and metabolism are an important consideration moving forward, these studies suggest that the results reported here fall within a dose range safe and relevant to translation into humans.

Children born with POE represent a growing population with poor long-term outcomes and limited treatment options, placing an increasing strain on healthcare and social support systems (3,6,7,68). Sodium butyrate is a cheap, accessible and safe postbiotic treatment widely available at most pharmacies as a dietary support. This, in combination with our findings, suggests NaB as a candidate for future clinical research, with potential to improve neurodevelopmental outcomes in this vulnerable group. Further research should further evaluate dosing and pharmacokinetic profiles during gestation and early development, while regulatory approval for prenatal use will also necessitate rigorous toxicological testing and evaluation of risk to benefits ratio to maternal and infant populations. More broadly this study provides compelling evidence linking gut-microbiome integrity to CNS development and cognition in rodent models of POE.

## Supporting information

Supplementary Statistics

Supplementary Results

## Acknowledgments

The authors acknowledge use of facilities in the Katharine Gaus Light Microscopy Facility at Mark Wainwright Analytical Centre for sample paraffin embedding and the Ramaciotti Centre for Genomics, UNSW for sequencing of the microbiome samples and Dovile Anderson from the Faculty of Pharmacy and Pharmaceutical Sciences, Monash University, Melbourne, Australia who ran the SCFA analysis. Experiments were performed on the UNSW Compute Cluster Katana (DOI: 10.26190/669X-A286). All timelines, schematics and explanatory figures were produced with Biorender.com. We would like to thank Melissa J. Bebbington for her invaluable assistance in teaching us immunofluorescence techniques, guiding us through the imaging process, and supporting our analysis using ImageJ.

## Data availability

Documentation detailing the allocation of offspring to experimental outcomes for each dam is available at http://hdl.handle.net/1959.4/106186.

