## Supplementary Methods for "Sodium Butyrate Rescues Neurodevelopmental Deficits Following Perinatal Methadone Exposure"

*Supplementary Methods i: Maternal Care Behaviors*

Behavioural definitions of maternal care are based on Myers et al. (1): Arched back nursing: when the dam was arched over her feeding pups with legs splayed; licking and grooming: dam licking or grooming any of her pups; blanket nursing: dam over the pups with no obvious arch in her back or extension of her legs; and passive nursing: the dam lying on her side with at least one pup nursing. Selfcare behavior was recorded when the dam was engaged in self-grooming, eating, drinking or exploration of her cage.

*Supplementary Methods ii: Trial Unique Non-Matching to Location Task Protocol*

The Trial Unique Non-Matching to Location (TUNL) task was performed in CANTAB boxes (Campden Instruments, Leicestershire, UK) composed of trapezoid-shaped chambers housed within sound attenuating boxes controlled by Whiskers Sever and ABET II software (2). Each box was equipped with a house light, a touch sensitive LCD screen covered with a mask to display 15 locations (5x3) and a magazine with pellet dispenser (45MG Dustless Precision Pellets, Able Scientific, AUS) on the opposite wall.

Pretraining, TUNL training, and testing were performed as previously described (3) with some modifications. Across pretraining stages, described in table i, rats were trained to touch one of the 15 locations illuminated on the screen to obtain a sucrose pellet paired with a tone. Incorrect responses – a touch to a non-illuminated location – resulted in a 5 second time-out where the house light was illuminated.

Following pretraining rats progressed to task training, where all sessions were 60 minutes or 100 trials, which ever elapsed first. Briefly, during task training the session was initiated by collection of a free pellet delivered to the magazine. One of the 15 available locations on the touch screen was illuminated, once the rat touched this location (S1) illumination ceased and the magazine was illuminated. The rat would then nose-poke the magazine, and two locations on the screen were illuminated, the initial location (S1) and a new location (S2). A correct response was recorded when the rat touched the new location (S2), a pellet was delivered to the illuminated magazine and a tone was played. After a 20 second ITI the next trial began with illumination of the magazine. An incorrect response was recorded if the rat nose-poked the initial location (S1) for a second time, and the response was followed by a 5 second time-out with the house light on, followed by the 20 second ITI. Over each session the illuminated locations varied in a pseudorandom random fashion. Locations with greater distance between them (i.e. “easy” separation 2, figure iC) are considered easier to differentiate than locations closer together (i.e. “hard” separation 0, figure 1A). The largest possible separations (i.e. separation 3, figure iD) were not used as they can be solved using strategies other than spatial and working memory, and no stacked combinations (i.e. Example stacked figure iE) were used as rats show a bias for the lower location (4).

Once 10 sessions of the TUNL task had been completed, rats were trained to a criterion of 75% correct responses on separation 2 (S2) trials for two consecutive days, or a total 20 sessions of TUNL (10 additional training days). Rats then progressed to the delay trials where an increasing temporal delay was introduced between nose poke of the initial location and illumination of the magazine. Two sessions each were run at 2, 4, 6 and 8 second delays.

The dependent variables considered in analysis of the TUNL task were accuracy (correct responses/correct + incorrect responses), premature responses, latency to collect reward and latency to make a correct response.

Supplementary Methods Table i. Summary of TUNL pretraining phases and criteria

| Phase | Phase description | Progression Criteria |
| --- | --- | --- |
| Habituation | 20 pellets in magazine | All pellets consumed within 30 minutes |
| Initial touch training | All grid windows lit together, rat touches any location while illuminated, tone is played, and 2 x pellets delivered | 30 minutes or 50 trials for one day |
| Must Touch training 1 | All grid windows lit together, rat touches any location while illuminated and receives 1 x food reward | 60 minutes or 100 trials for one day |
| Must Touch training 2 | Block of 2 x 2 windows are lit together to form a square. Unlit windows display a dark grey image. Rat touches any location while illuminated and receives 1 x food reward | 60 minutes or 100 trials for two consecutive days |
| Must Touch training 3 | One window is lit. Unlit windows display a dark grey image. Rat touches any location while illuminated and receives 1 x food reward | 60 minutes or 100 trials for two consecutive days |
| Must initiate training | A free delivery of food is made, and the tray light is turned on. The rat must nose poke and exit the magazine before a stimulus is displayed randomly on the screen. The rat must touch the stimulus to elicit tone/food response | 60 minutes or 90 trials for two consecutive days |
| Punish incorrect touches training | Same as must initiate training except incorrect touches lead to a TO period (5s) and no reward. | 60 minutes or 80% correct trials for two consecutive days |

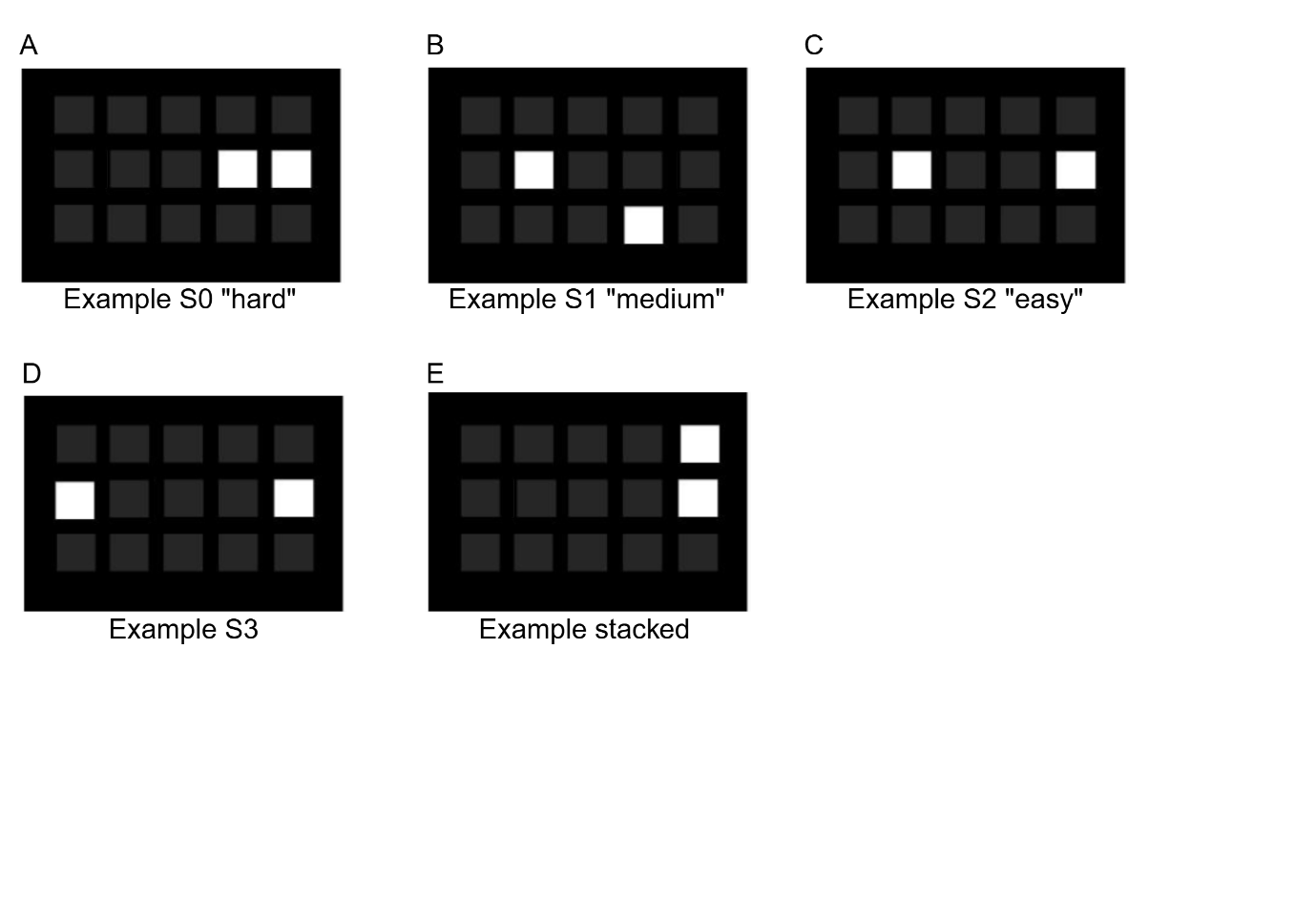

**Supplementary Methods Figure i: Representation of phase two stimulus locations in the TUNL task.** **A** represents a “hard” separation 0 (S0) trial, **B** a “medium” separation 1 (S1) trial and **C** an “easy” separation 2 (S2) trial used in the current study. **D** Represents a separation 3 (S3) trial and **E** a stacked configuration – these were not used as they do not rely on spatial memory alone.

*Supplementary Methods iii: Five-Choice Serial Reaction Time Task Protocol*

The 5-CSRTT was performed in a modified operant chamber (MED Associates, VA, US) individually housed in sound-attenuating cabinets. The left wall of the chamber was curved with five apertures, each equipped with an LED light and infrared detectors. On the opposite wall, a magazine with a ceiling light was connected to a pellet dispenser. Rats progressed through the 5-CSRTT according to the criteria and conditions at each stage outlined in table ii. Each session lasted 30 minutes or 100 trials, whichever elapsed first, and rats were required to meet criteria in two consecutive sessions or a maximum number of days before progressing to the next stage.

Each session began with the illumination of the magazine and delivery of a pellet. Once the rat had collected the pellet a delay (inter trial interval; ITI) began. Thereafter one of the 5 nose-poke apertures was illuminated in a pseudorandom fashion, with the stimulus duration (SD) varying based on the stage of the task (summarized in table ii). If the rat nose-poked the illuminated aperture within a specified time (limited hold) a correct response was recorded, and a pellet delivered. The next trial would commence once that pellet was collected. If the rat made a response during the ITI (premature response), made a response to the wrong aperture during the limited hold (incorrect response), or failed to nose-poke the correct aperture during the limited hold (omission) a 5 second time out occurred where the house light turned on. The next trial commenced after the time out. Any entries into the nose-poke apertures made after a correct response but before collection of the reward from the magazine were recorded as perseverative responses but did not result in a time-out. Once rats had progressed through all the training stages, they underwent one multiple stimulus duration (MSD) session, a single session at stage 7 the following day and then one multiple ITI (MITI) session.

Measures of performance were accuracy (correct responses/correct + incorrect responses), omissions (omissions/total number of trials), premature responses and perseverative responses.

Supplementary Methods Table ii. Summary of 5-CSRTT phases and progression criteria

| Phase | Stimulus duration (seconds) | Inter-trial interval (seconds) | Limited hold (seconds) | Progression criteria |
| --- | --- | --- | --- | --- |
| Habituation | - | - | - | Pellets consumed from each nose-poke aperture (2) and magazine (5) within 20 minutes |
| 1 | 30 | 2 | 30 | ≥ 30 correct trials |
| 2 | 20 | 2 | 20 | ≥ 30 correct trials |
| 3 | 10 | 5 | 10 | ≥ 50 correct trials |
| 4 | 5 | 5 | 5 | ≥ 50 correct trials, 75% accuracy and  < 20% omissions |
| 5 | 2.5 | 5 | 5 | ≥ 50 correct trials, 75% accuracy and  < 20% omissions  < 20% omissions |
| 6 | 1.25 | 5 | 5 | ≥ 50 correct trials and 75% accuracy  < 20% omissions  < 20% omissions |
| 7 | 1 | 5 | 5 | ≥ 50 correct trials and 75% accuracy  < 20% omissions or a maximum of 12 days |
| MSD | 0.25, 0.5, 1, 1.25, 2.5 | 5 | 5 | One session |
| MITI | 1 | 1, 3, 5, 7 | 5 | One session |

*Supplementary Methods iv: Tissue Processing and Immunofluorescence (P21)*

Mounted sections were de-paraffinized in Histochoice Clearing Agent (Merck, Macquarie Park, CAS# H2779) and rehydrated with graded ethanol baths for 10 minutes (100%, 95% 70%, 50%, deionized water). Sections then underwent antigen retrieval through immersion in Tris buffer (pH 9; Promega, Alexandria, CAS# V6232) while heated for 3 minutes in a microwave. Following three, 5-minute washes in PBS, sections were pre-incubated in 10% normal donkey serum (NDS) for 1-hour, followed by overnight incubation with a primary antibody in 2% NDS solution. The primary anti-bodies rabbit polyclonal anti-onized calcium binding adaptor molecule 1 (Iba1) antibody (1:100, 109-19741, Novachem, Victoria, Australia), mouse monoclonal anti-glial fibrillary acidic protein (GFAP) antibody (1:200; MAB360; Merck Millipore, Darmstadt, Germany) and rabbit monoclonal anti-myelin binding protein (MBP) antibody (1:500, M3821, Merck Millipore). The following day, sections were washed 3x 5-minute in PBS, and then incubated with a secondary antibody in PBS, (all dilutions 1:250; 488-conjugated donkey anti-mouse; cy3-conjugated donkey anti-rabbit; Thermo Fisher, MA, USA), for 30 minutes. After three 5-minute washes in PBS, sections were cover-slipped with DAPI fluoroshield mounting medium (F6057, Sigma-Aldrich) for nuclear staining.

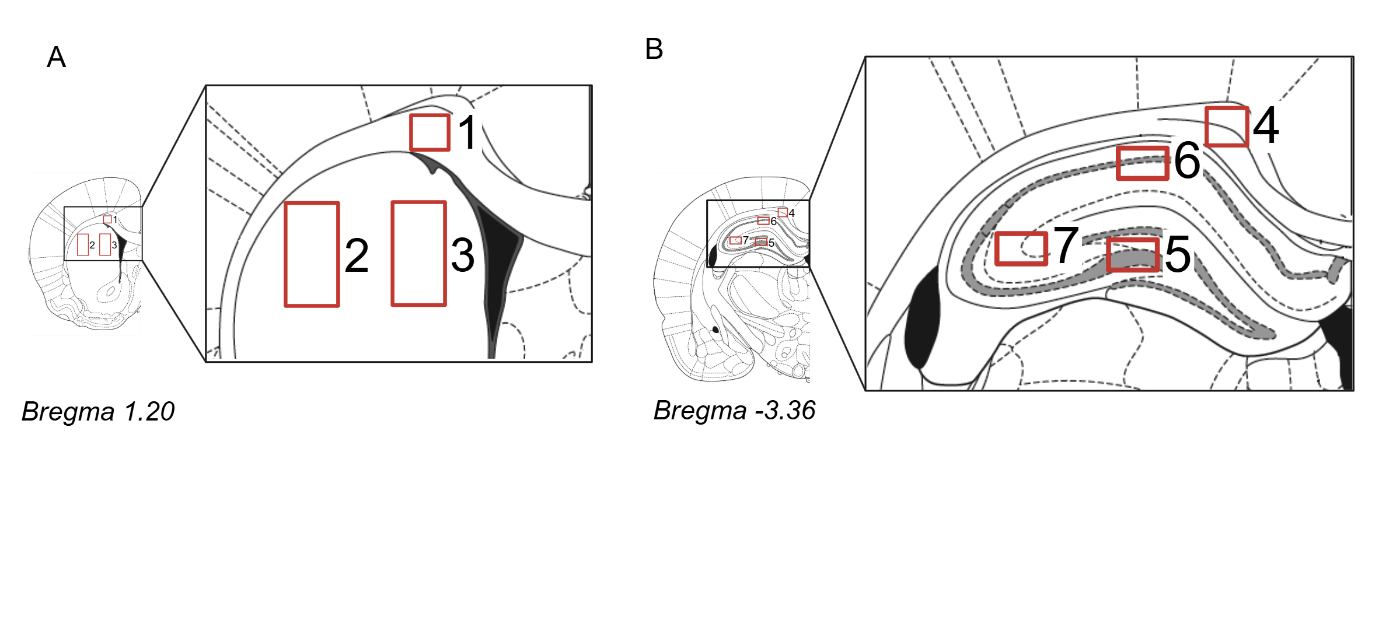

**Supplementary Methods Figure ii: Regions of interest for immunofluorescence**. **A** Striatum including 1. anterior corpus callosum, 2 dorsal lateral striatum and 3. dorsal medial striatum. **B** Dorsal hippocampus including 4. posterior corpus callosum, 5. dentate gyrus, 6. CA1 and 7. CA3 of the hippocampus. Images are adapted from the Paxinos & Watson Rat Brain Atlas (5).

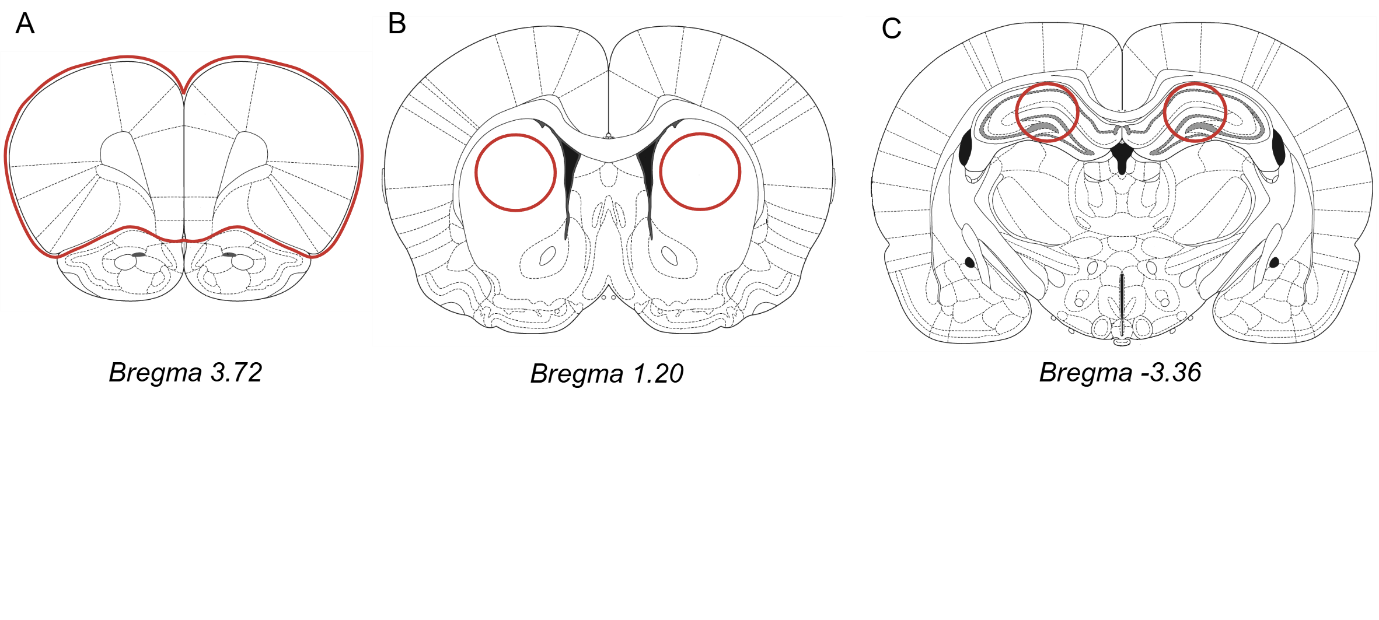

**Supplementary Methods Figure iii: Representative regions collected for RT-qPCR analysis from P21 animals.** Areas outlined in red were collected for analysis in **A** the prefrontal cortex, **B** the striatum, and **C** the dorsal hippocampus.

Supplementary Methods Table iii: TaqMan assays used to target genes of interest

| Gene name | Assay ID |
| --- | --- |
| Immune |  |
| IL-10 | Rn01483988_g1 |
| IL-6 | Rn01410330_m1 |
| IL-1B | Rn00580432_m1 |
| TNFα | Rn99999017_m1 |
| CD68 | Rn01495634_g1 |
| Myelination |  |
| MBP | Rn01399619_m1 |
| SOX 10 | Rn00569909_m1 |
| Myrf | Rn01454573_m1 |
| Housekeeping genes |  |
| UBC | Rn01789812_g1 |
| GapDH | RN01775763_g1 |

*Supplementary Methods v: Cytokine Multiplex*

At P7 and 21 blood was collected from the perfusion circuit, held at room temperature for 30 mins, centrifuged at 1,000 x g for 10 mins and then the supernatant (serum) collected. Serum was used to assess levels of 23 cytokines and chemokines using a magnetic bead immunoassay (Bio-Plex Pro™ Rat Cytokine 23-Plex Assay #12005641) in duplicates at a 1:4 dilution following manufacturer’s instructions. The analytes considered are summarized in table iv. The median fluorescent intensity for the 23 analytes was obtained for all standards. Median fluorescence intensity was used to allow for analysis of low abundance analytes (6). Samples with a coefficient of variation greater than 25% were removed from analysis in that assay.

Supplementary Methods Table iv: Analytes considered in cytokine multiplex’s analyzing serum collected on postnatal day 7 and postnatal day 21

| Analyte | Function |
| --- | --- |
| Interleukin-1α (IL-1α) | Proinflammatory cytokine |
| IL-1β | Proinflammatory cytokine |
| IL-18 | Proinflammatory cytokine |
| IL-17a | Proinflammatory cytokine |
| Macrophage inflammatory protein-1α (MIP-1α) | Proinflammatory chemokine |
| MIP-3a | Proinflammatory chemokine |
| Monocyte chemoattractant protein-1 (MCP-1) | Proinflammatory chemokine |
| Interferon gamma (IFN-γ) | Proinflammatory cytokine |
| Tumour necrosis factor- α (TNF-α) | Proinflammatory cytokine |
| IL-6 | Pro and anti-inflammatory cytokine |
| IL-4 | Anti-inflammatory cytokine |
| IL-10 | Anti-inflammatory cytokine |
| IL-13 | Anti-inflammatory cytokine |
| IL-2 | T cell growth factor |
| IL-5 | Th2 cytokine |
| IL-7 | T cell growth factor |
| Regulated on activation normal T cell expressed and secreted (RANTES) | Chemokine |
| Growth-related oncogene/keratinocyte derived chemokine (GRO-KC) | Chemokine |
| Granulocyte-macrophage colony-stimulating factor (GM-CSF) | Growth factor/colony stimulating factors |
| Macrophage colony-stimulating factor (mCSF) | Growth factor/colony stimulating factors |
| Vascular endothelial growth factor (VEGF) | Growth factor/colony stimulating factors |
| IL-12p70 | Adaptive immunity |

*Supplementary Methods vi: In Vivo Intestinal Permeability*

*Animals*

*Dams:* 39 female and 4 male Sprague Dawley rats were purchased from Animal Resource Centre, Perth, Australia. All animals were maintained on a 12-hour day/night light cycle (lights on at 0700h) in a temperature- and humidity-controlled colony room (22°C±1) with *ad libitum* access to irradiated rat and mouse diets SF00-100 (Specialty Feeds, Western Australia, Australia) and water unless otherwise specified. Following a minimum of one week habituation to the facility breeding began, where two females were housed with one male. Following pairing females were swabbed twice daily (between 0800h and 1000h, and 1600h and 1800h) until sperm was sighted or a week elapsed. The day sperm was sighted was designated GD0, and dams were separated and single housed until giving birth, as previously described (7). All dams gave birth within 24 hours of GD22 (designated postnatal day 0 [P0]).

*Pups:* On P0 litter sizes were standardized to 10 pups (5 female, 5 male where possible).

*Drugs*

Methadone hydrochloride (National Institute of Measurement, North Ryde, Australia) was administered identically to the procedure described in the main text.

1.5% w/v sodium butyrate (Sigma-Aldrich, Missouri, USA) was administered via drinking water (refreshed every 48-72 hours) across GD10-P10 to mimic human prenatal development. This concentration is known to be tolerated by rats and result in increased fecal levels of butyric acid (8), with lower levels administered during gestation and lactation known to result in increased plasma levels of butyrate in pups (9).

**Procedures**

*Dams*

On GD9 dams were randomly allocated to one of three experimental conditions: sham surgery/water (Sham/Veh), methadone pump/water (Met/Veh), methadone pump/sodium butyrate (Met/NaB). When pregnancy was not detected by GD9, dams were allocated to Sham and underwent sham surgery on P3 only. Final dam group allocations were Sham/Veh=11, Met/Veh=17, Met/NaB=9.

*Pups*

One male and one female pup from each litter underwent assessment in vivo assessment of gut permeability as described in the main text.

*Supplementary Method vii: Morphological Analysis of Microglia in the Dentate Gyrus at P21*

To assess the impact of drug exposure on microglial morphology further analysis was performed on the images acquired from the dentate gyrus (Supplementary Methods Figure ii). Images were analyzed in ImageJ (National Institutes of Health, Bethesda, US) using the Analyze Skeleton (10) and Sholl analysis (11) functions. The protocol was based on a previously published work (12). All image processing was performed blinded to condition.

For whole image skeleton analysis, images of the dentate gyrus were opened in ImageJ and a manual cell count performed. The images were then duplicated and a macro was run to apply the functions Unsharp Mask, Despeckle, transform to 8-bit, threshold “Triangle” applied, Binary close, Skeletonize, and Analyze Skeleton. The output generated was then sorted by the number of branches from highest to lowest, and only the number of identified skeletons that corresponded to the number of cells manually counted where retained. The values of these skeletons for each outcome from an image were averaged and used for analysis.

For individual microglia skeleton analysis a macro was run on a further duplicate to apply the functions Unsharp Mask, Despeckle, transform to 8-bit, threshold “Triangle” applied and Binary close. Seven microglia from this processed image were then selected randomly and, using the original image as a reference, back ground noise manually removed and microglia processes manually joined. On these individual images the processes Skeletonize and Analyze Skeleton were then run. The output for each outcome from the cells from an image was averaged and used for analysis.

For Sholl analysis the same individual images were used, prior to the application of Skeletonize and Analyze Skeleton. A radius was drawn from the center of the cell body and Sholl analysis run. The output for each outcome from the cells from an image was averaged and used for analysis.
