## Supplementary Results for "Sodium Butyrate Rescues Neurodevelopmental Deficits Following Perinatal Methadone Exposure"

**Supplementary Results**
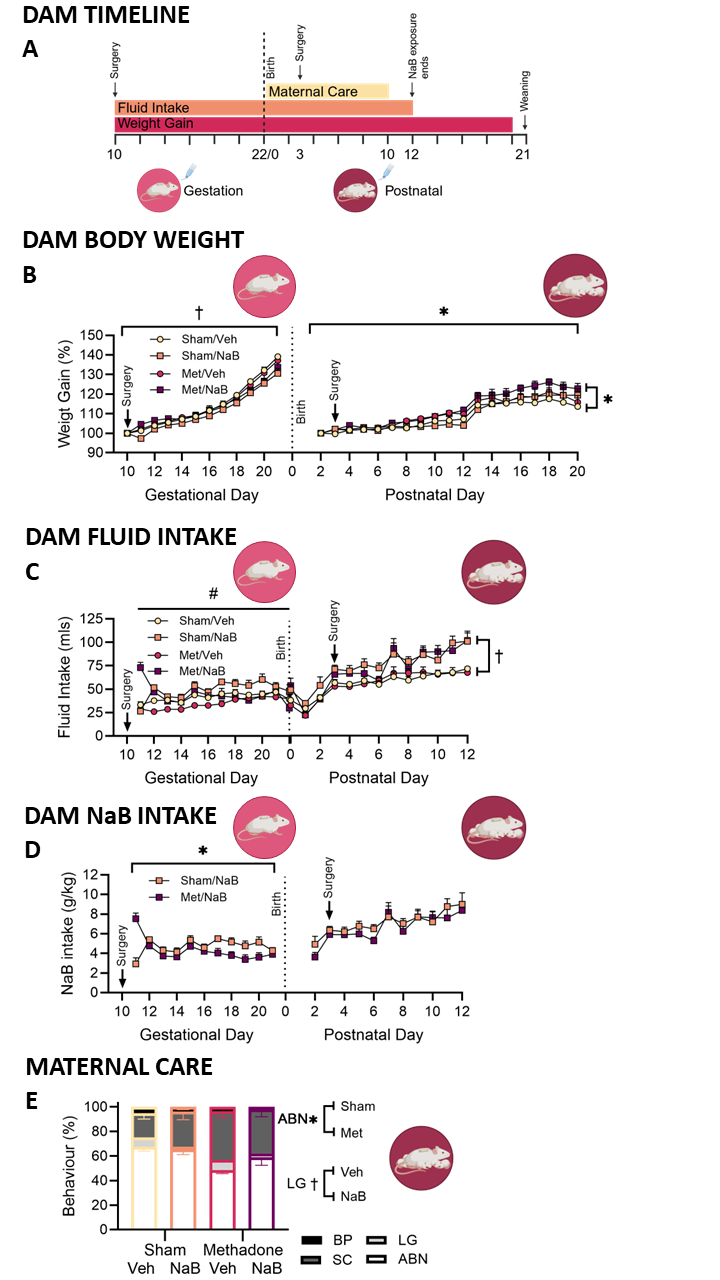

**Supplementary Results Figure i: Dams gestational and postnatal body weight, fluid intake and maternal care. A** Timeline of dam drug exposure across the gestational and postnatal period, **B** Dam relative weight gain (% baseline body weight) recorded daily from gestational day 10 (the start of drug exposure), until the day prior to birth, and following birth from postnatal day 2 until postnatal day 20. **C** Dam fluid intake across the gestational and postnatal period recorded daily from gestational day 10 until postnatal day 12 when NaB exposure ceased. **D** Dam NaB intake g/kg across the gestational and postnatal period recorded daily from gestational day 10 until the day prior to birth, and following birth from postnatal day 2 until postnatal day until postnatal day 12 when NaB exposure ceased, as rats were not weight across the expected birth period. Data represented as group mean + SEM with 6/7 rats/group. **E** Between postnatal day 1 and 10 dams maternal care behavior was recorded daily. Data is represented as percentage of observations in which dams were seen to be engaged in each maternal care behavior, arched back nursing (ABN), licking and grooming (LG), selfcare behavior (SC) and blanket and passive nursing (BP). Maternal care behavior did not correlate with any adult behavioral outcomes. Data represented as group mean + SEM with Sham/Veh n=6, Sham/NaB n = 6, Met/Veh n=6, Met/NaB n=. Asterisk, obelisk and hashtag indicate a significant effect of pump, fluid or pump by fluid interaction respectively.

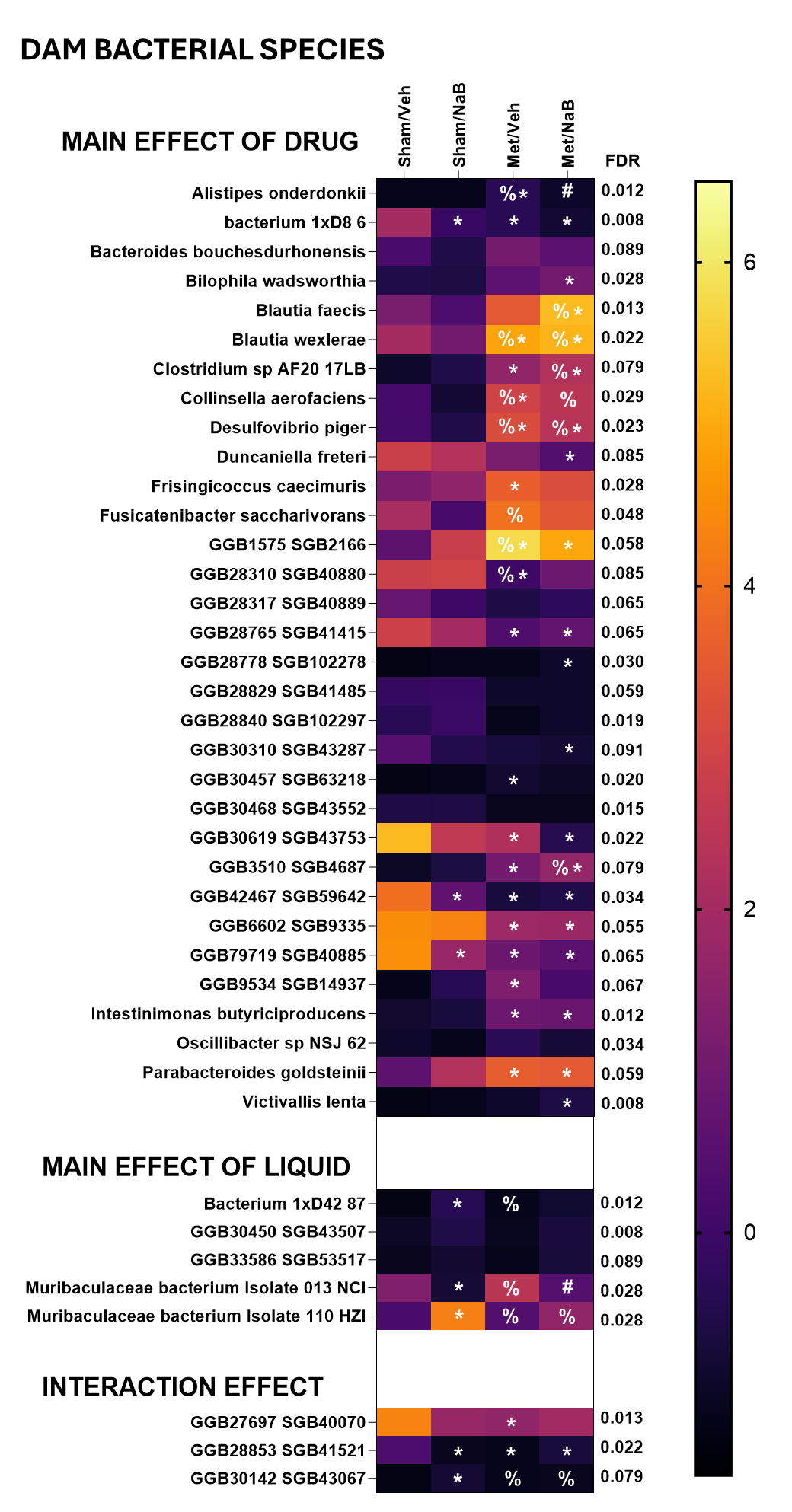

**Supplementary Results Figure ii: Microbial species and functions altered by methadone, sodium butyrate or their interaction in dam faecal microbiota.** Heatmaps representing the relative abundance of bacterial species significantly altered by treatments. Data as averaged central logarithmic ratios. Significant FDR-corrected main and interaction effects are reported to the right of the heat maps. Sham/Veh n=6, Sham/NaB n = 6, Met/Veh n=6, Met/NaB n=7. Tukey-adjusted post-hoc comparisons *p<0.05 relative to Sham/Veh; #p<0.05 relative to Met/Veh; % p<0.05 relative to Sham/NaB.

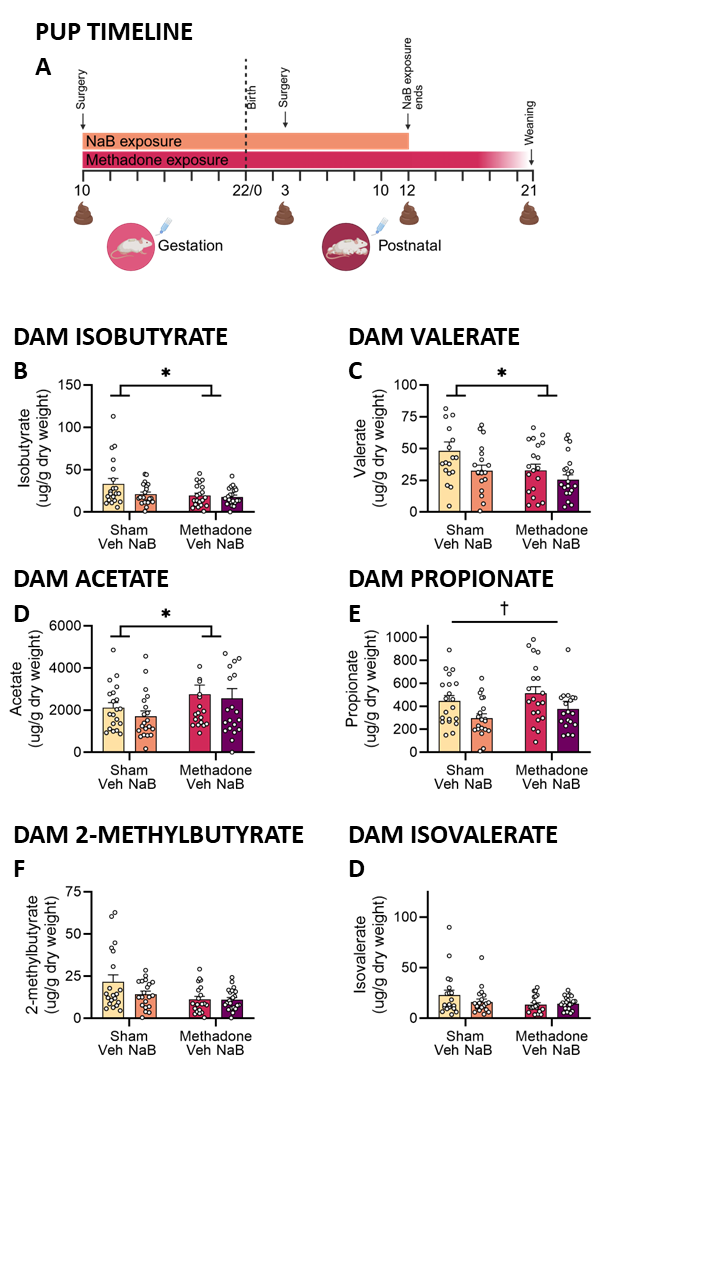
 **Supplementary Results Figure iii: Dams fecal short chain fatty acid concentration. A** Timeline of dam drug exposure and fecal collection across the gestational and postnatal period. Concentration of **B** Isobutyrate, **C** Valerate, **D** Acetate, **E** 2-propionate, **F** 2-methyburtyrate, and **G** Isovalerate adjusted for sample volume and moisture content (ug.g dry weight). Data represented as group mean + SEM with Sham/Veh n=6, Sham/NaB n = 6, Met/Veh n=6, Met/NaB n=7. Asterisk, obelisk indicate a significant effect of pump or fluid respectively.

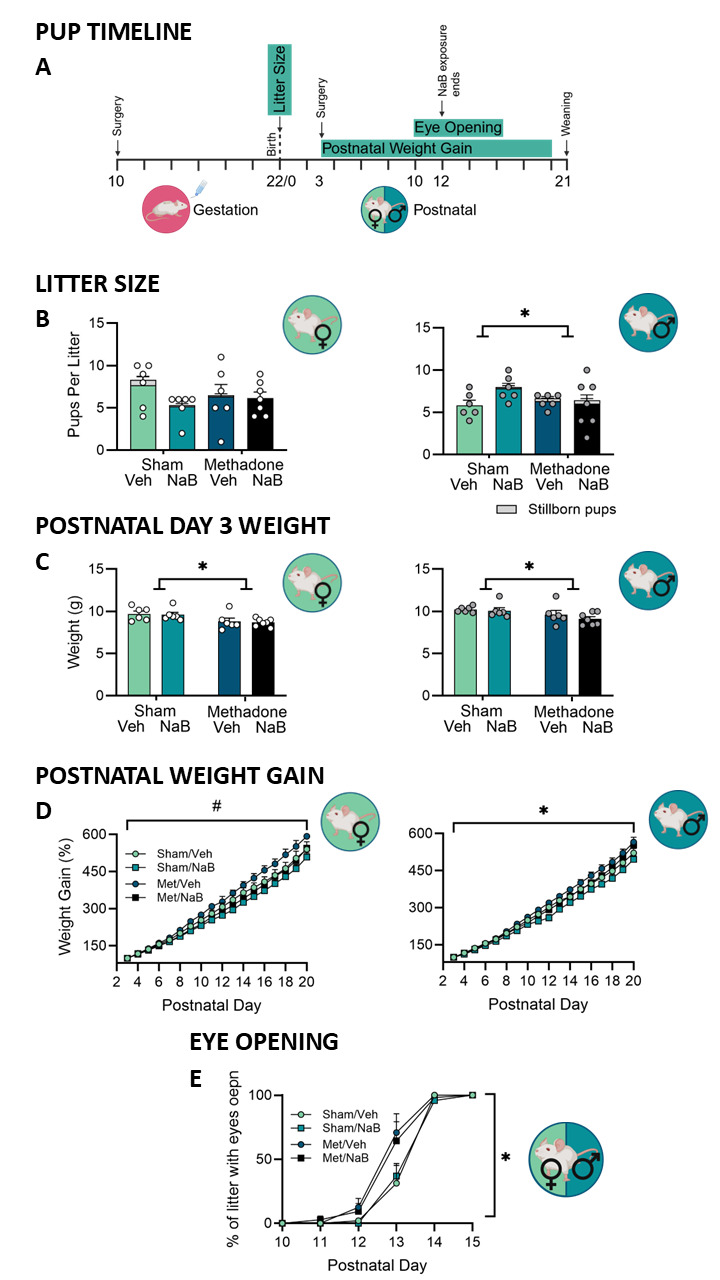

**Supplementary Results Figure iv: Pups litter size and number of still born pups, weight on postnatal day 3, weight gain prior to weaning, and day of eye opening. A** Pup timeline. **B** Litter size and number of still born pups recorded on P0. **C** Pups were first weighed on postnatal day 3, shown here as the average weight of male and female pups in a litter. Data points represent group mean **±** SEM with 6/7 female (white points) and male (grey points) pups/group. **D** Pups were then weighed daily until postnatal day 20. To account for initial differences in weight at P3 relative weight gain (% baseline body weight) was calculated from P3, with the average female and male weight used from each litter. **E** The percentage of each litter with both of their eyes open was recorded daily from postnatal day 10 to 15. Data points represent group mean **±** SEM with female (white points, Sham/Veh n=6, Sham/NaB n = 6, Met/Veh n=6, Met/NaB n=7) and male (grey points, Sham/Veh n=6, Sham/NaB n = 6, Met/Veh n=6, Met/NaB n=7) pups. Asterisk and hashtag indicate a significant main effect of pump or pump by fluid interaction respectively.

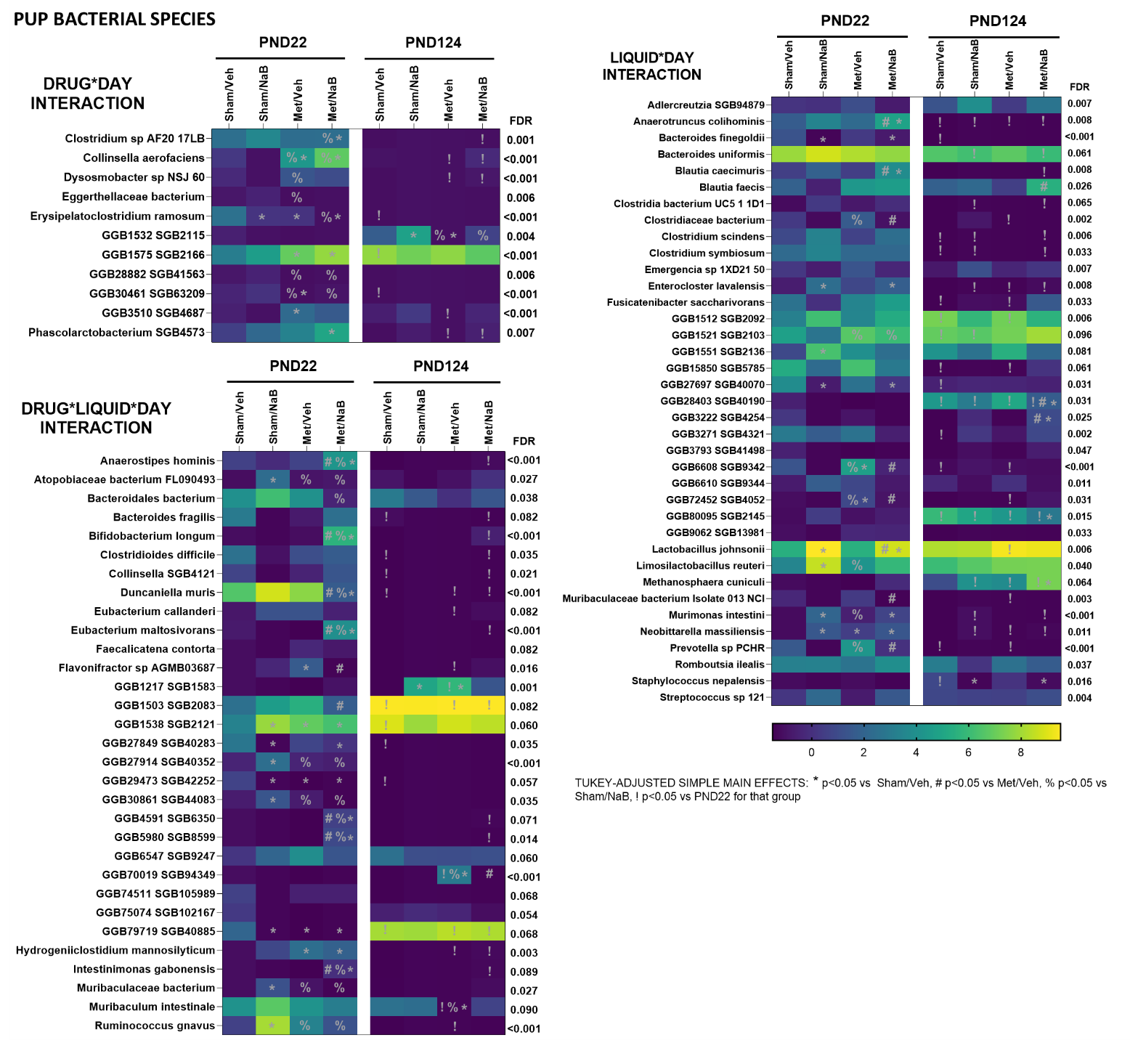

**Supplementary Results Figure v: Microbial species altered by methadone, sodium butyrate or their interaction across development in pup faecal microbiota.** Heatmaps representing the relative abundance of bacterial species significantly altered by treatments Data as averaged central logarithmic ratios. Significant FDR-corrected main and interaction effects are reported to the right of the heat maps. Tukey-adjusted post-hoc comparisons P21 females Sham/Veh n=6, Sham/NaB n = 6, Met/Veh n=6, Met/NaB n=7 and males Sham/Veh n=6, Sham/NaB n = 6, Met/Veh n=6, Met/NaB n=7. At P124 there were 2-4 pups/sex/litter included in analysis from Sham/Veh n=6, Sham/NaB n = 6, Met/Veh n=6, Met/NaB n=7 litters, with litter controlled for using a linear mixed effects model approach. Specific numbers for each group are as follows – females Sham/Veh n=10, Sham/NaB n=5, Met/Veh n=8, Met/NaB n=9, males, Sham/Veh n=9, Sham/NaB n=5, Met/Veh n=8, Met/NaB n=12. *p<0.05 relative to Sham/Veh; #p<0.05 relative to Met/Veh; % p<0.05 relative to Sham/NaB; !p<0.05 relative to PND22 for that group.

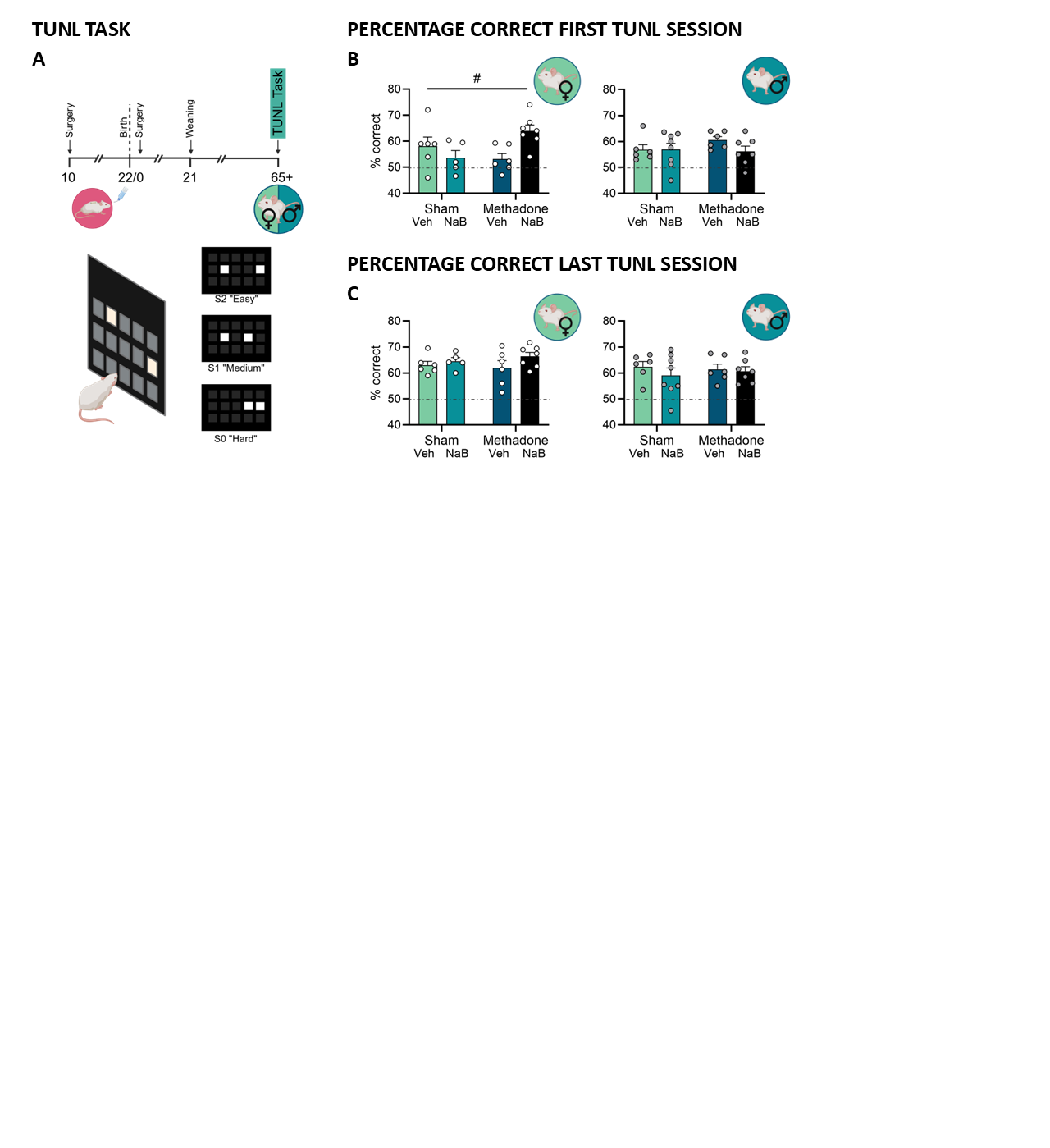

**Supplementary Results Figure vi: Performance during TUNL task acquisition. A** Schematic of the TUNL task, which included one female and one male pup (P65) from each litter. **B** Percentage of correct trials on the first session of the TUNL task, where the spatial component of the task was introduced. **C** Percentage of correct trials on the last session of the .5s delay sessions, before progression to 2, 4, 6 and 8 second delay. Data points represent group mean **±** SEM with female (white points Sham/Veh n=6, Sham/NaB n = 5, Met/Veh n=6, Met/NaB n=7) and male (grey points Sham/Veh n=6, Sham/NaB n = 8, Met/Veh n=6, Met/NaB n=7) pups. The dashed horizontal line represents chance performance, with no preference for the novel location. Hashtag indicates a significant pump by fluid interaction.

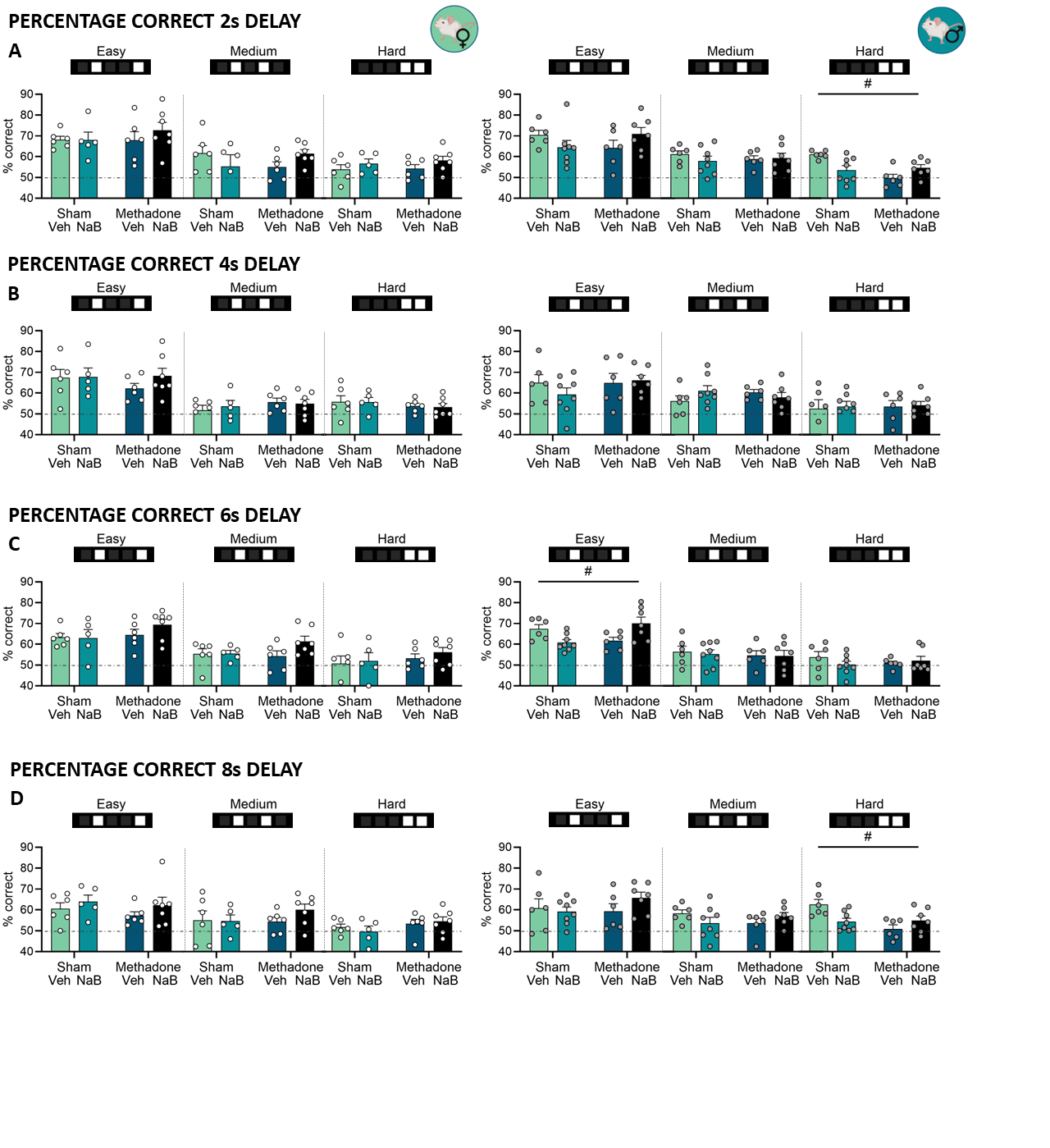

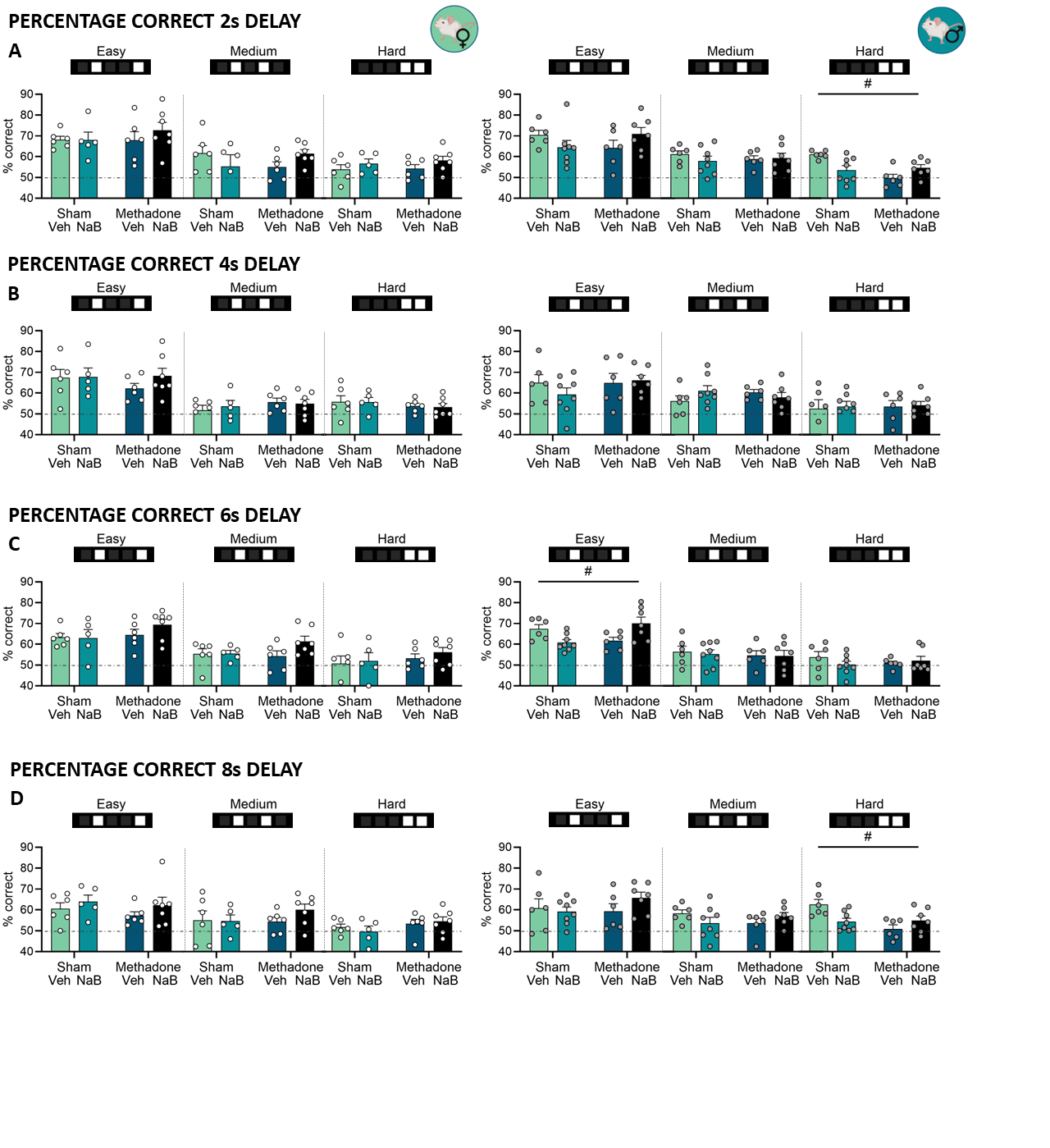

**Supplementary Results Figure vii: Performance on the TUNL task separated by spatial difficulty and temporal delay.** Percentage of correct responses in **A** 2 second, **B** 4 second, **C** 6 second and **D** 8 second delay sessions, separated by spatial difficulty. Data points represent group mean **±** SEM with female (white points Sham/Veh n=6, Sham/NaB n = 5, Met/Veh n=6, Met/NaB n=7) and male (grey points Sham/Veh n=6, Sham/NaB n = 8, Met/Veh n=6, Met/NaB n=7) pups. The dashed horizontal line represents chance performance, with no preference for the novel location. Hashtag indicate a significant pump by fluid interaction.

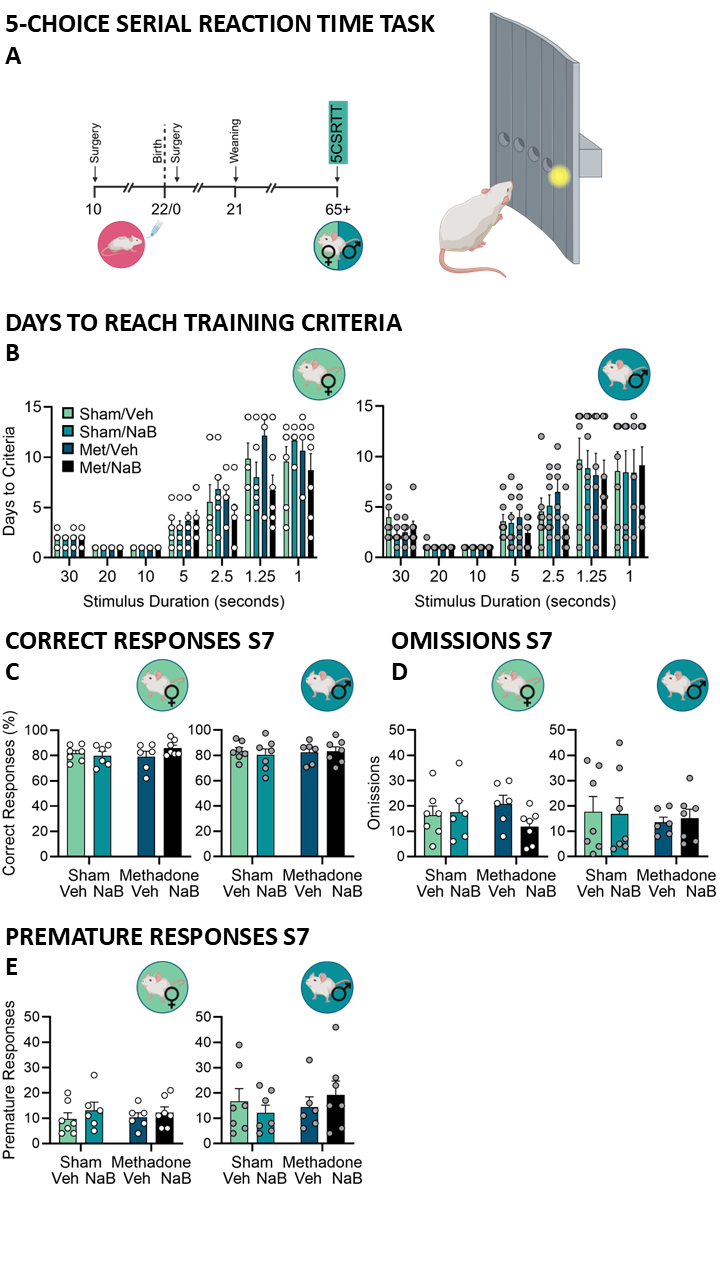

**Supplementary Results Figure viii: Acquisition of the 5-choice serial reaction time task. A** Schematic of the 5-CSRTT task which included one female and one male pup (P65) from each litter. **B** Number of days taken to reach criteria at each stage. Data points represent group mean **±** SEM with female (white points Sham/Veh n=7, Sham/NaB n = 6, Met/Veh n=6, Met/NaB n=7) and male (grey points Sham/Veh n=7, Sham/NaB n = 7, Met/Veh n=6, Met/NaB n=7) pups.

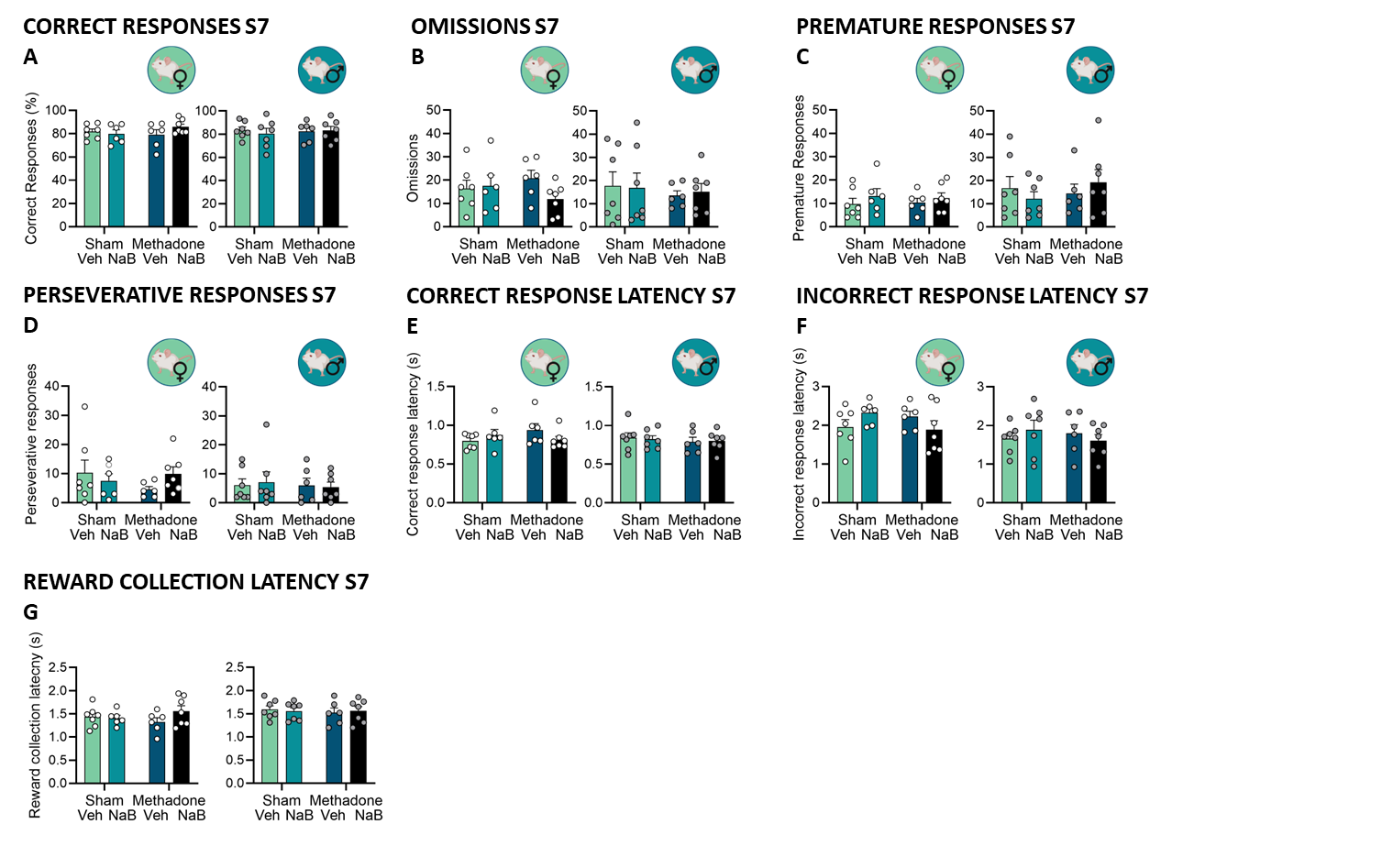

**Supplementary Results Figure ix: Performance in stage 7 of the 5-choice serial reaction time task. A** Percentage of correct responses in the final stage 7 session. **B** Number of omissions in the final stage 7 session. **C** Number of premature responses in the final stage 7 session. **D** Number of perseverative responses in the final stage 7 session. **E** Correct response latency in the final stage 7 session. **F** Incorrect response latency in the final stage 7 session. **G** reward collection latency in the final stage 7 session. Data points represent group mean **±** SEM with female (white points Sham/Veh n=7, Sham/NaB n = 6, Met/Veh n=6, Met/NaB n=7) and male (grey points Sham/Veh n=7, Sham/NaB n = 7, Met/Veh n=6, Met/NaB n=7) pups.

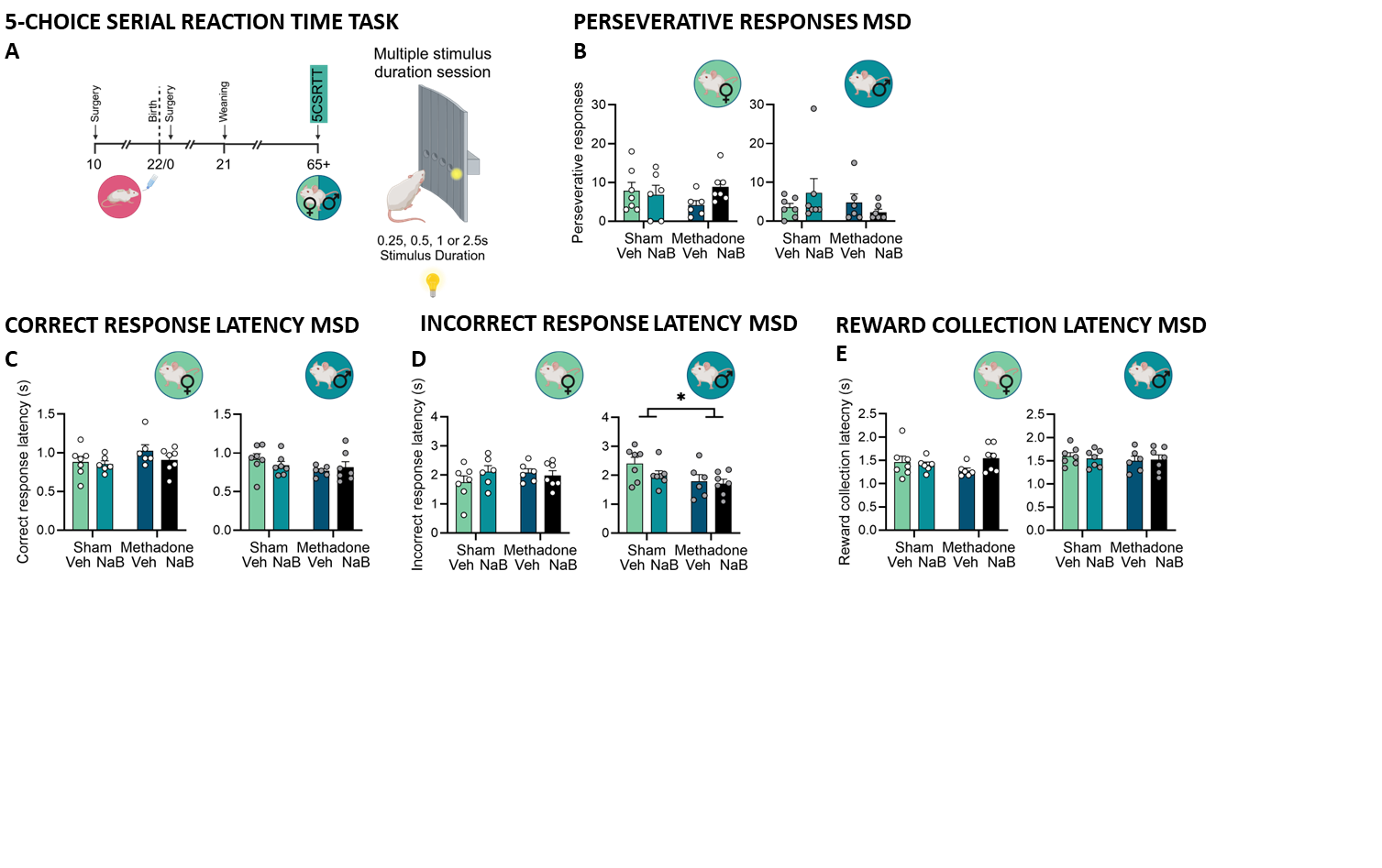

**Supplementary Results Figure x: MSD performance in the 5-choice serial reaction time task. A** Schematic of the 5-CSRTT task which included one female and one male pup (P65) from each litter. **B** Number of perseverative responses in the MSD session. **C** Correct response latency in the MSD session. **D** Incorrect response latency in the MSD session. **E** reward collection latency in the MSD session. Data points represent group mean **±** SEM with female (white points Sham/Veh n=7, Sham/NaB n = 6, Met/Veh n=6, Met/NaB n=7) and male (grey points Sham/Veh n=7, Sham/NaB n = 7, Met/Veh n=6, Met/NaB n=7) pups. Asterisks indicate a significant main effect of pump.

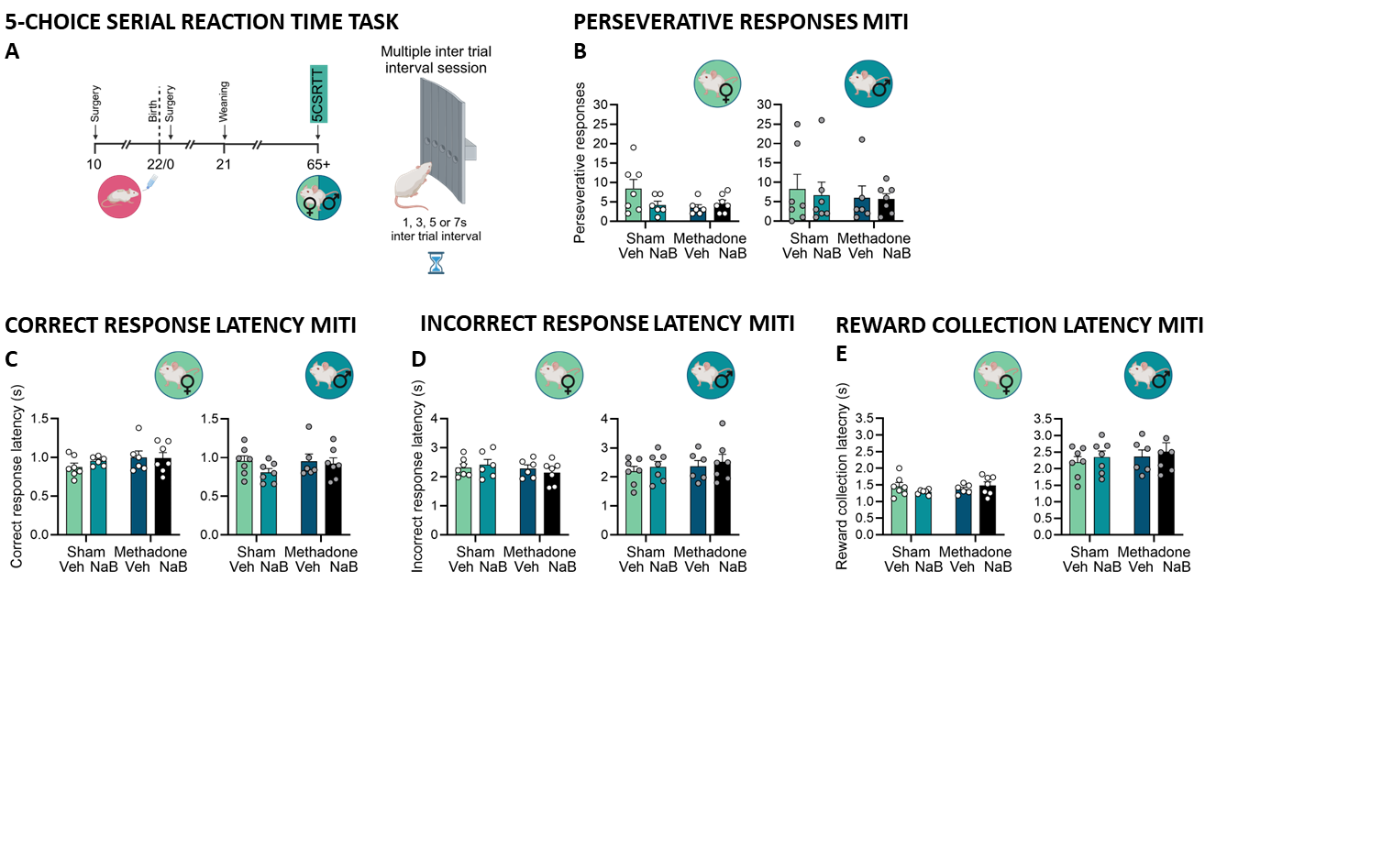

**Supplementary Results Figure xi: MITI performance in the 5-choice serial reaction time task. A** Schematic of the 5-CSRTT task which included one female and one male pup (P65) from each litter. **B** Number of perseverative responses in the MITI session. **C** Correct response latency in the MITI session. **D** Incorrect response latency in the MITI session. **E** reward collection latency in the MITI session. Data points represent group mean **±** SEM with female (white points Sham/Veh n=7, Sham/NaB n = 6, Met/Veh n=6, Met/NaB n=7) and male (grey points Sham/Veh n=7, Sham/NaB n = 7, Met/Veh n=6, Met/NaB n=7) pups.

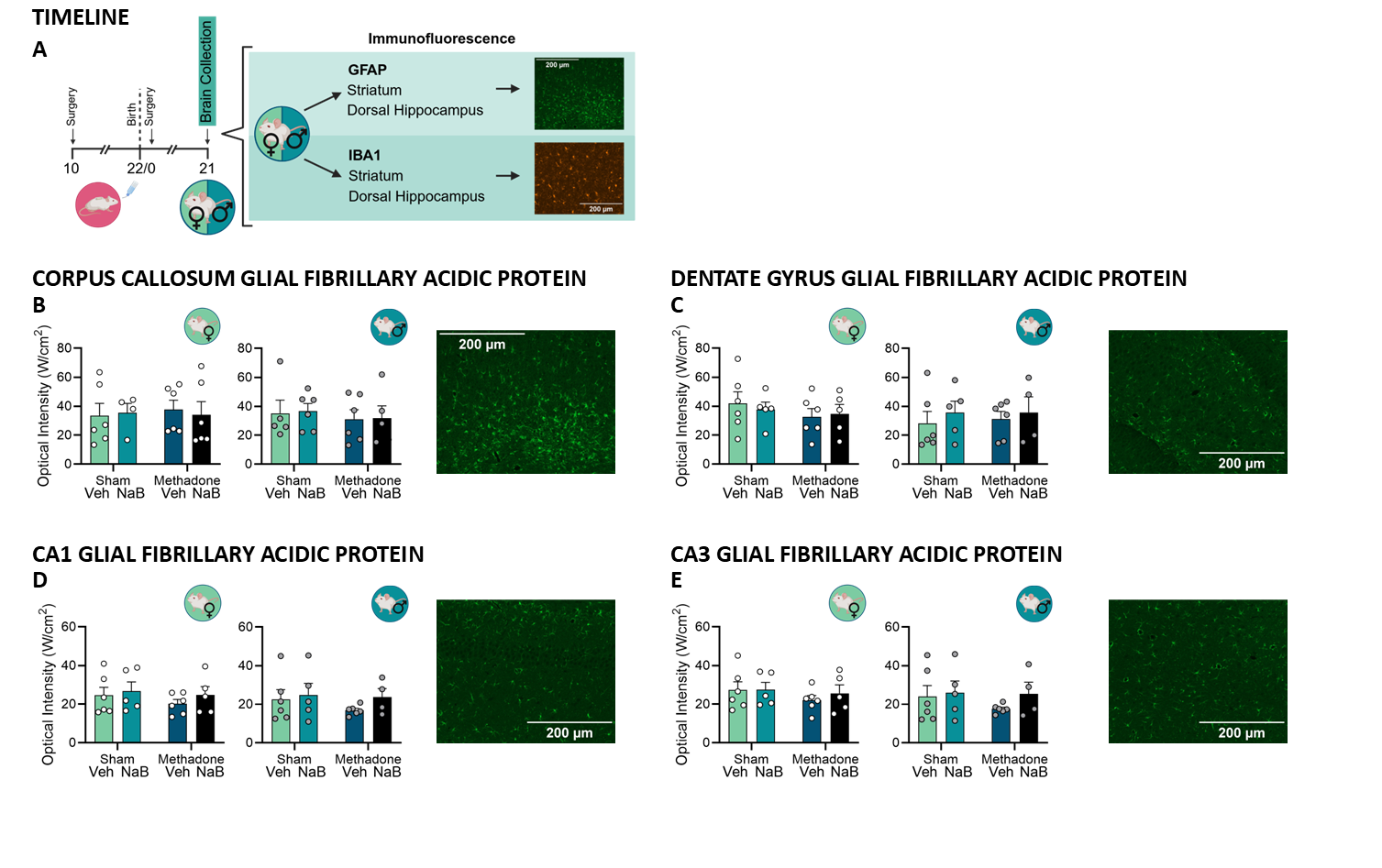

**Supplementary Results Figure xii: Staining for Glial Fibrillary Acidic Protein (GFAP) at postnatal day 21. A** Timeline of brain collection and areas stained. The optical intensity of staining for GFAP in **B** the corpus callosum at the level of the striatum, **C** the dentate gyrus, **D** the CA1, and **E** the CA3. Data points represent group mean **±** SEM with female (white points Sham/Veh n=6, Sham/NaB n = 5, Met/Veh n=6, Met/NaB corpus callosum n=6, hippocampus n=5) and male (grey points Sham/Veh n=6, Sham/NaB n=6, hippocampus n=5, Met/Veh n=6, Met/NaB corpus callosum n=6, hippocampus n=5) pups.

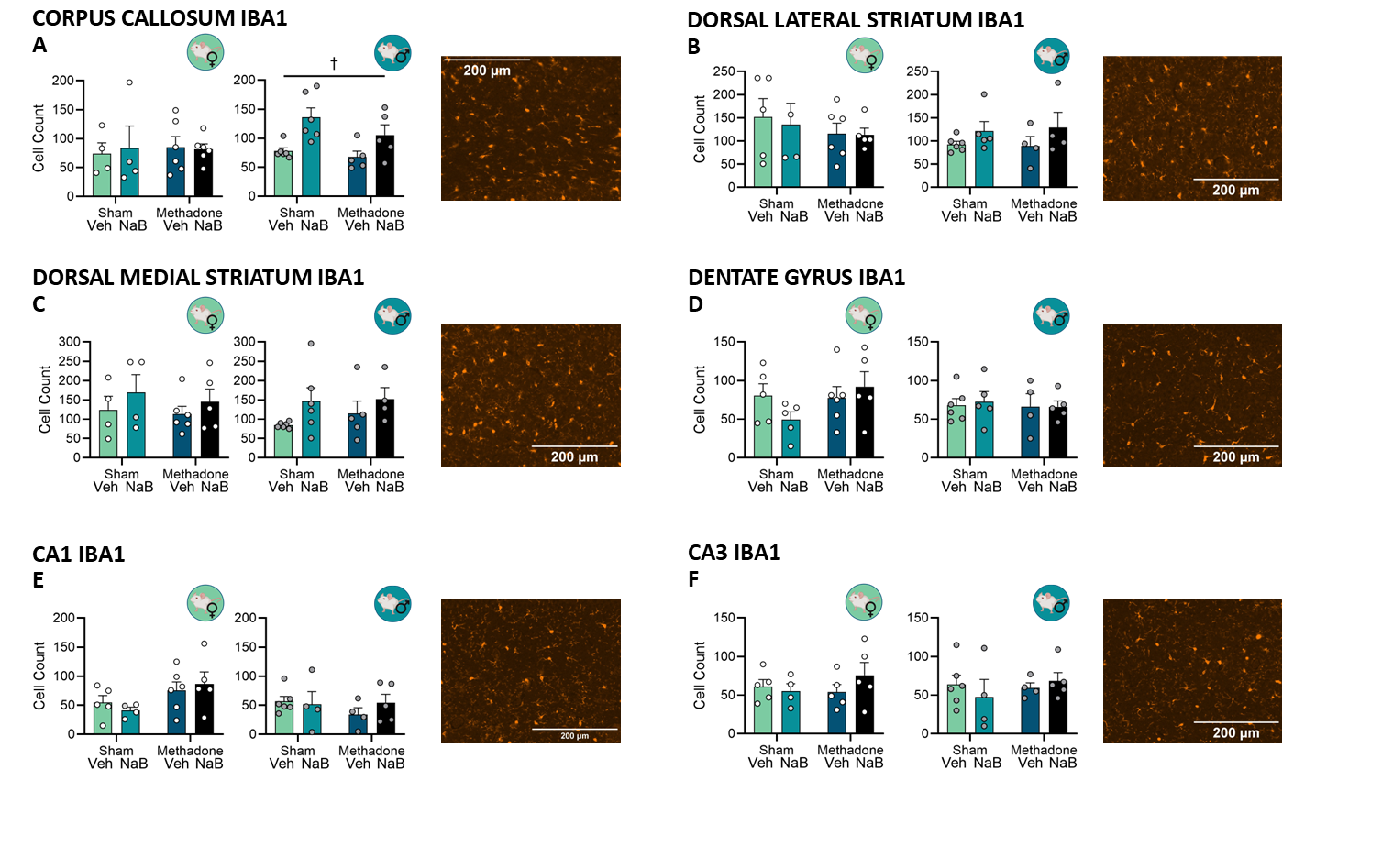

**Supplementary Results Figure xiii: Staining for Ionized calcium binding adaptor molecule 1 (IBA1) at postnatal day 21.** The number of IBA1 positive cells stained in **A** the corpus callosum at the level of the striatum, **B** the dorsal lateral striatum, **C** the dorsal medial striatum, **D** the dentate gyrus, **E** the CA1, and **F** the CA3. Data points represent group mean **±** SEM with female (white points Sham/Veh striatum n=4, hippocampus n= 5, Sham/NaB striatum n=4, hippocampus n= 5, Met/Veh n=6, Met/NaB striatum n=6, hippocampus n=5) and male (grey points Sham/Veh n=6, Sham/NaB striatum n=6, hippocampus n=5, Met/Veh striatum n=5, hippocampus n=4, Met/NaB n=5) pups. Obelisk indicates a significant main effect of fluid.

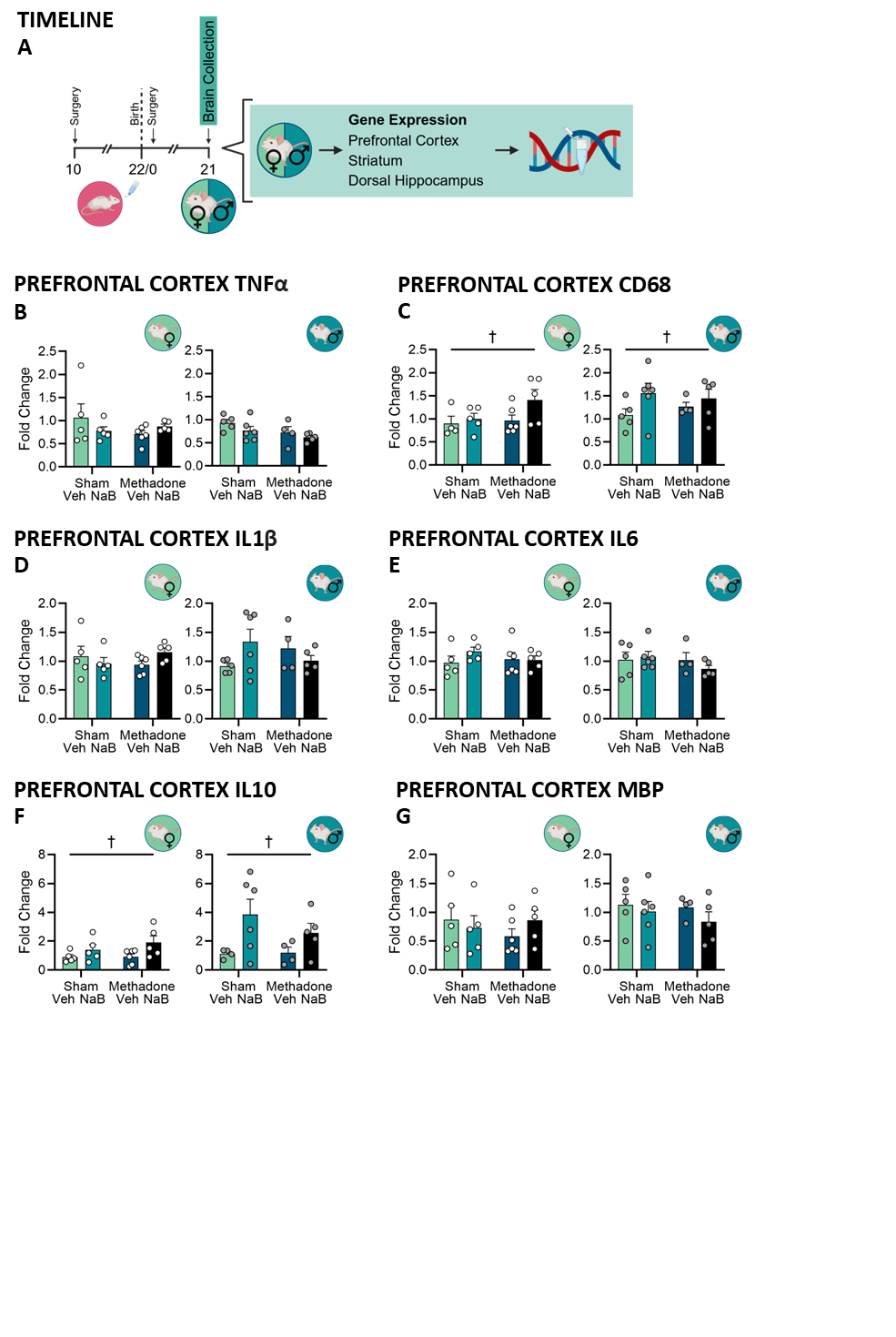

**Supplementary Results Figure xiv: Gene expression in the prefrontal cortex (PFC) on postnatal day 21. A** Timeline of brain collection and regions processed for RT qPCR. **B** The relative expression in the PFC of **B** TNFα, **C** CD68, **D** IL1β, **E** IL6, **F** IL10 and **G** MBP. Data points represent group mean **±** SEM with female (white points Sham/Veh n=5, Sham/NaB n=5, Met/Veh n=6, Met/NaB n=5) and male (grey points Sham/Veh n=5, Sham/NaB n=6, Met/Veh n=4, Met/NaB n=5) pups. Obelisk indicates a significant main effect of fluid.

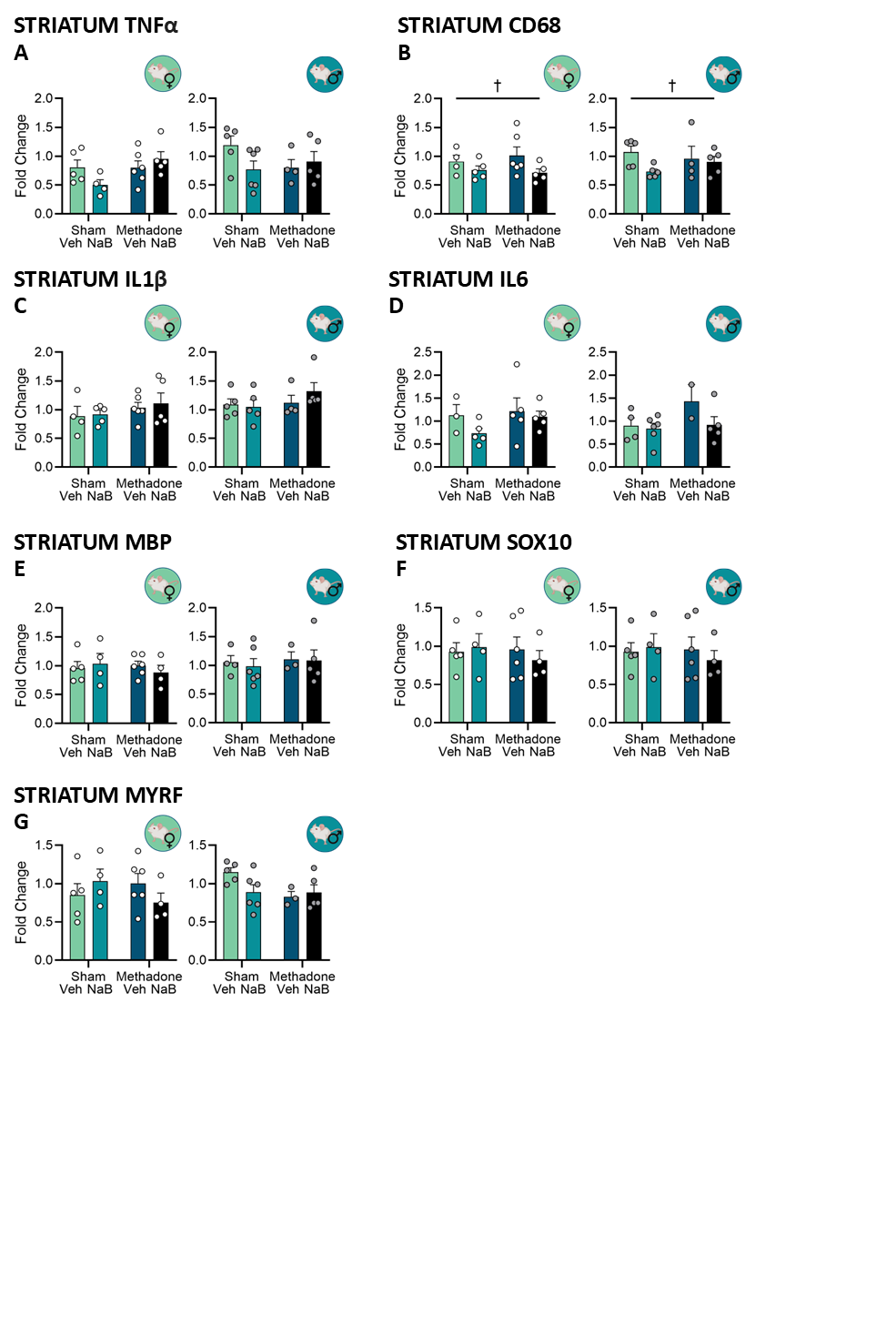

**Supplementary Results Figure xv: Gene expression in the striatum on postnatal day 21.** The relative expression in the striatum of **A** TNFα, **B** CD68, **C** IL1β, **D** IL6, **E** MBP, **F** SOX10, and **G** MYRF Data points represent group mean **±** SEM with female (white points Sham/Veh n=5, Sham/NaB n=5, Met/Veh n=6, Met/NaB n=5) and male (grey points Sham/Veh n=5, Sham/NaB n=6, Met/Veh n=4, Met/NaB n=5) pups=. Hanging obelisk indicates a significant main effect of fluid.

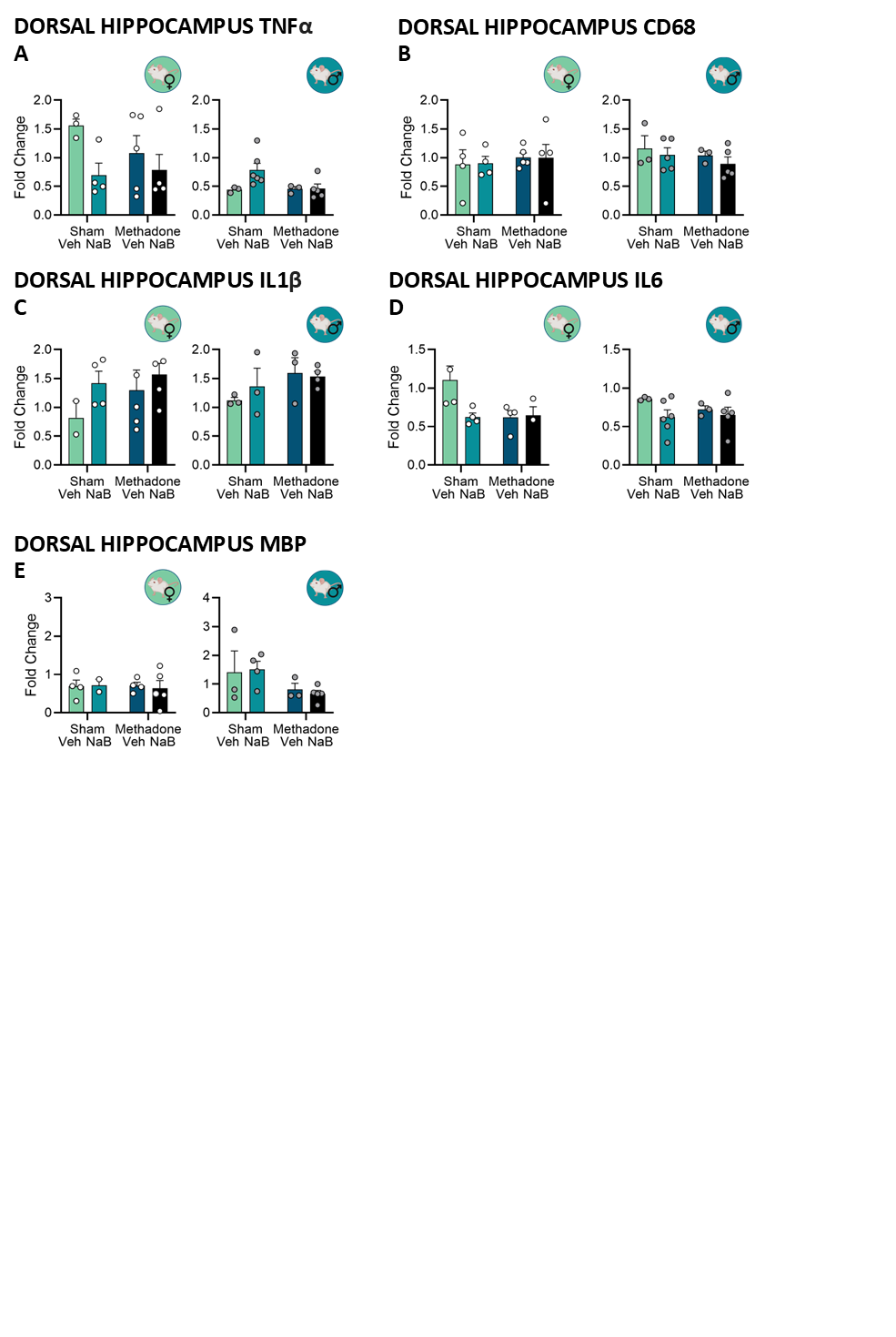

**Supplementary Results Figure xvi: Gene expression in the dorsal hippocampus (dHPC) on postnatal day 21.** Relative expression of **A** TNFα, **B** CD68, **C** IL1β, **D** IL6, **E** MBP, **F** SOX10 and **G** MYRF in dorsal hippocampus (dHPC). Data points represent group mean **±** SEM with female (white points Sham/Veh n=2/4, Sham/NaB n=4, Met/Veh n=5, Met/NaB n=5) and male (grey points Sham/Veh n=3, Sham/NaB n=4-6, Met/Veh n=3, Met/NaB n=5) pups.

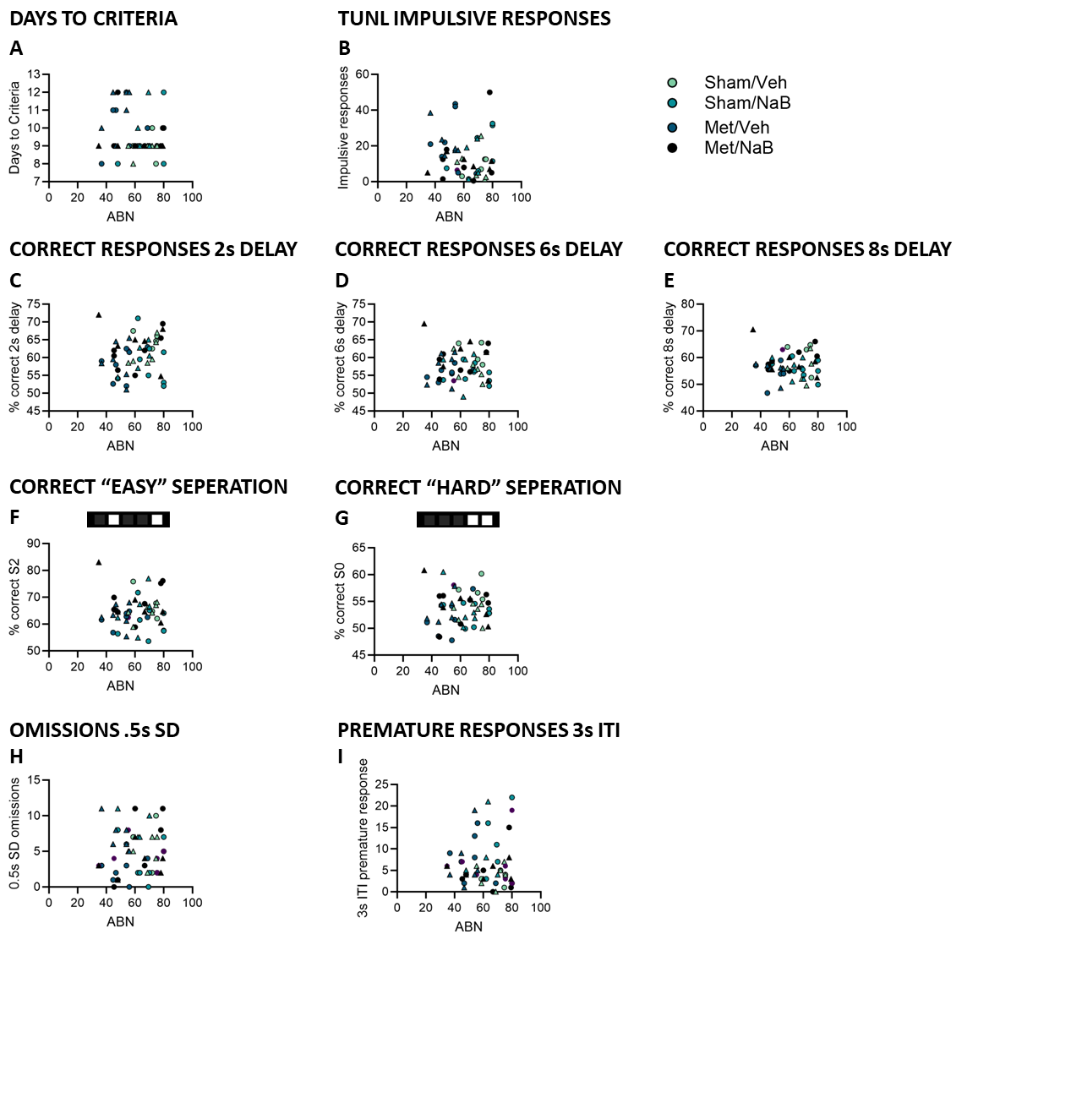

**Supplementary Results Figure xvii: Analysis of correlations between adult behavior and arch back nursing (ABN) from dam.** Graphs show the correlation between ABN and **A** days to criteria on the TUNL task, **B** impulsive responses in the TUNL task, **C** correct responses in the TUNL task at a 2 seconds (s) delay, **D** correct responses in the TUNL task at a 6s delay **E** correct responses in the TUNL task at an 8s delay, **F** in TUNL task sessions with “easy” (S2) separation trials, **G** in TUNL task sessions with “hard” (Ss) separation trials, **H** omissions during .5s trials on the multiple stimulus duration session of the 5CSRTT and **I** premature responses on 3s trials in the multiple inter trial interval session of the 5CSRTT. Data points represent group mean **±** SEM with female (triangle TUNL Sham/Veh n=6, Sham/NaB n=5, Met/Veh n=6, Met/NaB n=7, 5CSRTT Sham/Veh n=7, Sham/NaB n=6, Met/Veh n=6, Met/NaB n=7) and male (circle TUNL Sham/Veh n=6, Sham/NaB n=8, Met/Veh n=6, Met/NaB n=7, 5CSRTT Sham/Veh n=7, Sham/NaB n=6, Met/Veh n=6, Met/NaB n=7) pups.

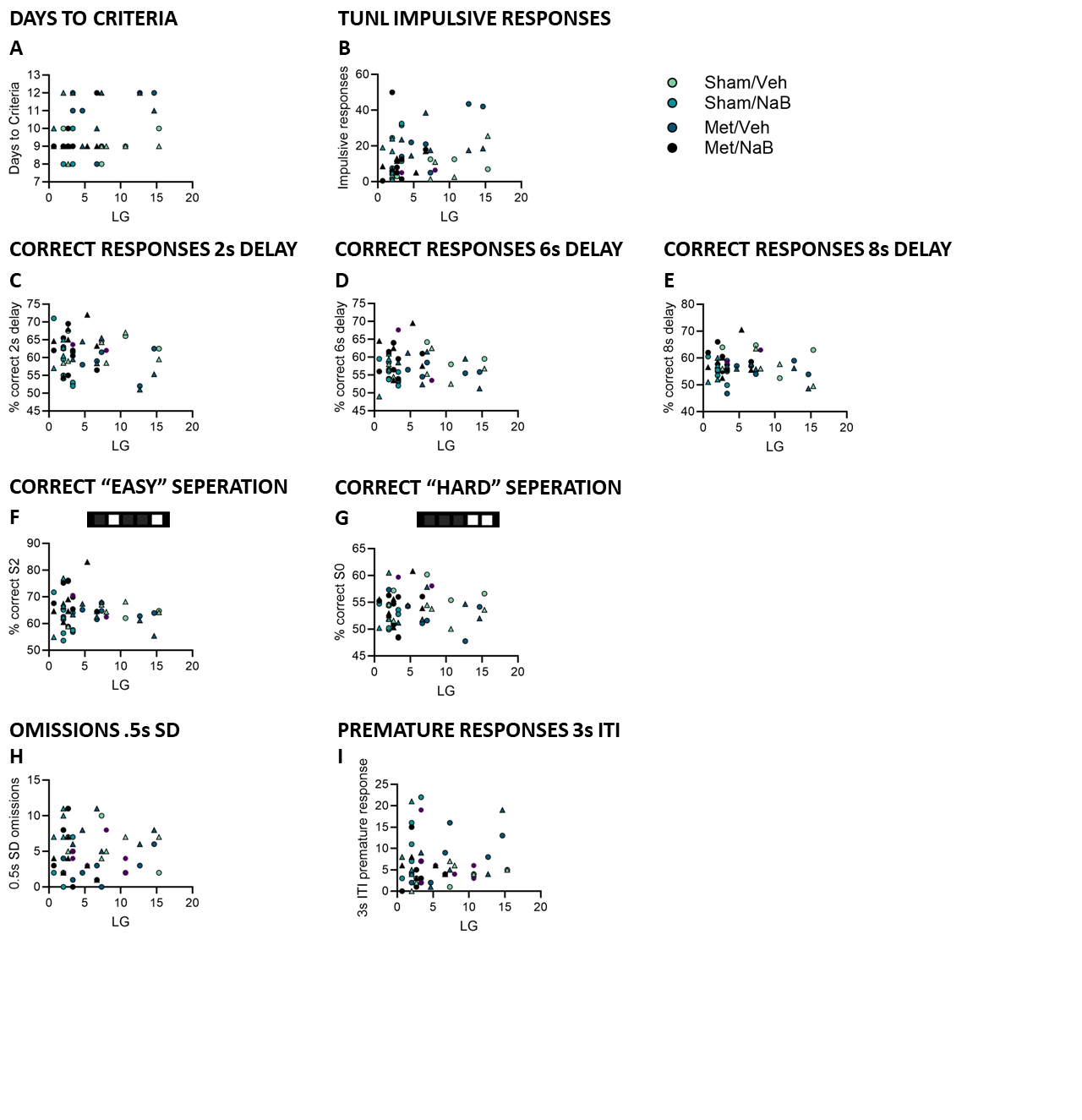

**Supplementary Results Figure xviii: Analysis of correlations between adult behavior and licking and grooming (LG) from dam.** Graphs show the correlation between LG and **A** days to criteria on the TUNL task, **B** impulsive responses in the TUNL task, **C** correct responses in the TUNL task at a 2 seconds (s) delay, **D** correct responses in the TUNL task at a 6s delay **E** correct responses in the TUNL task at an 8s delay, **F** in TUNL task sessions with “easy” (S2) separation trials, **G** in TUNL task sessions with “hard” (Ss) separation trials, **H** omissions during .5s trials on the multiple stimulus duration session of the 5CSRTT and **I** premature responses on 3s trials in the multiple inter trial interval session of the 5CSRTT. Data points represent group mean **±** SEM with female (triangle TUNL Sham/Veh n=6, Sham/NaB n=5, Met/Veh n=6, Met/NaB n=7, 5CSRTT Sham/Veh n=7, Sham/NaB n=6, Met/Veh n=6, Met/NaB n=7) and male (circle TUNL Sham/Veh n=6, Sham/NaB n=8, Met/Veh n=6, Met/NaB n=7, 5CSRTT Sham/Veh n=7, Sham/NaB n=6, Met/Veh n=6, Met/NaB n=7) pups/group.

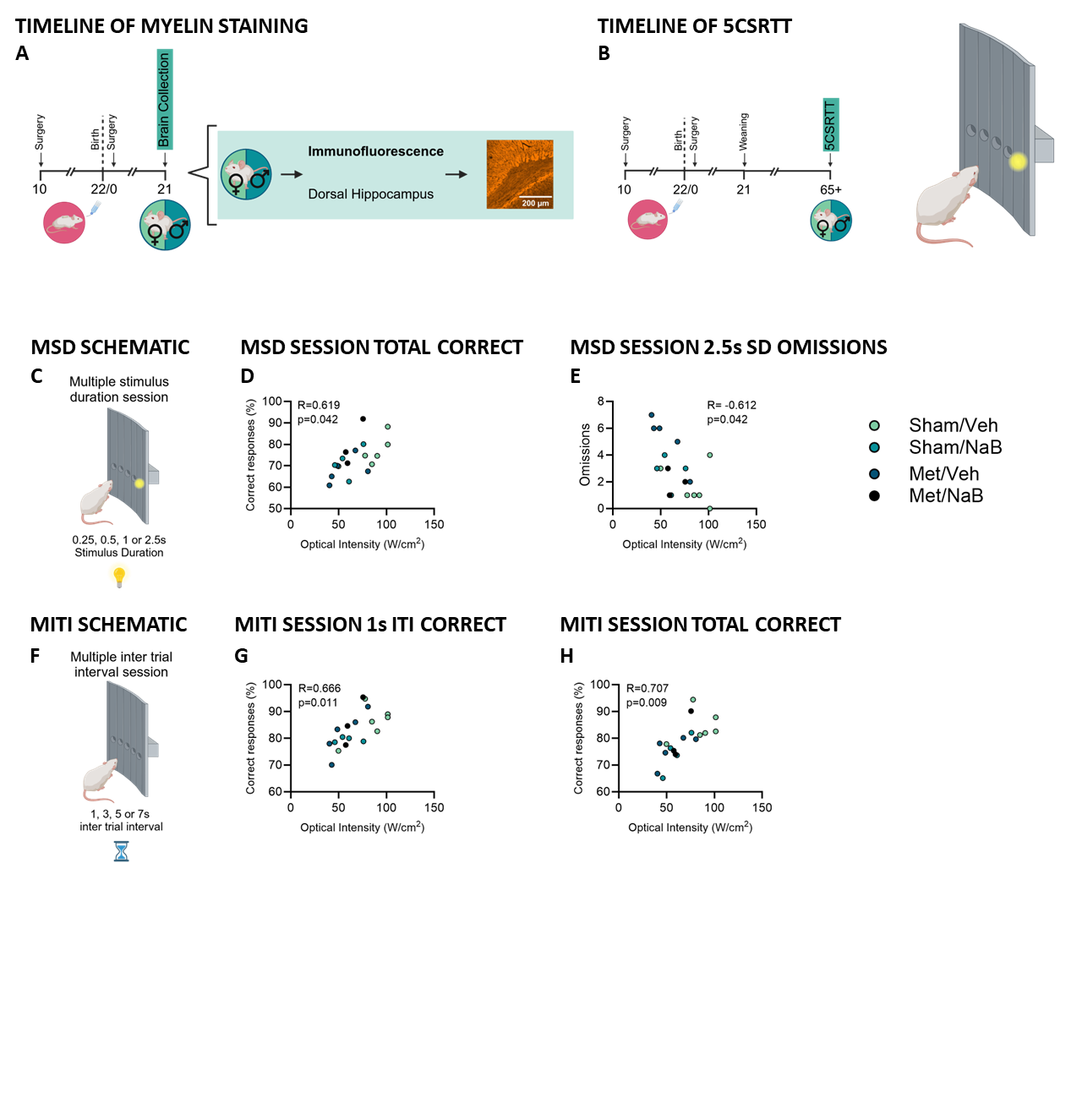

**Supplementary Results Figure xxiii: Exploratory correlational analysis of myelin basic protein staining in the corpus callosum at P21 and adult behavior performance between female siblings. A** Timeline of myelin basic protein (MBP) staining **B** Timeline of 5 choice serial reaction time task (5CSRTT) **C** Schematic of the multiple stimulus durations (MSD) session of the 5CSRTT **D** correlation between MBP staining in the corpus callosum and the percentage of correct responses in the MSD session of the 5CSRTT in females **E** correlation between MBP staining in the corpus callosum and number of omissions at 2.5s SD in the MSD session of the 5CSRTT in females **F** Schematic of the multiple inter trial interval (ITI) session of the 5CSRTT **G** correlation between MBP staining in the corpus callosum and the percentage of correct responses during 1s ITI in the MITI session of the 5CSRTT in females **H** correlation between MBP staining in the corpus callosum and the overall percentage of correct responses during the MITI session of the 5CSRTT in females. Data points represent individual rats, color coded for group with Sham/Veh n=6, Sham/NaB n=2, Met/Veh n=4, Met/NaB n=6, females.

| Supplementary Results Table i: Cytokine expression in pups’ serum at postnatal day 7 | | | | | | | | | | | | | | | | | |
| --- | --- | --- | --- | --- | --- | --- | --- | --- | --- | --- | --- | --- | --- | --- | --- | --- | --- |
|  | Sham/Veh Mean (SEM) | | | | Met/Veh Mean (SEM) | | | | Sham/NaB Mean (SEM) | | | | Met/NaB Mean (SEM) | | | |  |
|  | Male (n=6) | | Females (n=4) | | Male (n=5) | | Females (n=5) | | Male (n=5) | | Females (n=5) | | Male (n=5) | | Females (n=5) | | *p-*value |
| IL-13 | 1.008 | (0.115) | 1.177 | (0.042) | 0.988 | (0.133) | 1.026 | (0.161) | 1.208 | (0.033) | 1.226 | (0.023) | 1.135 | (0.027) | 1.135 | (0.028) | † .046 |
| IL-10 | 1.051 | (0.077) | 1.325 | (0.068) | 0.964 | (0.145) | 1.11 | (0.075) | 1.228 | (0.036) | 1.178 | (0.049) | 1.176 | (0.071) | 1.176 | (0.072) | † males .034 $ .034 |
| IL-1A | 1.187 | (0.183) | 1.392 | (0.082) | 1.45 | (0.065) | 1.252 | (0.082) | 1.462 | (0.076) | 1.508 | (0.096) | 1.38 | (0.061) | 1.38 | (0.062) | all > .05 |
| IL-2 | 1.166 | (0.034) | 1.26 | (0.015) | 1.225 | (0.015) | 1.154 | (0.044) | 1.236 | (0.016) | 1.218 | (0.047) | 1.232 | (0.045) | 1.232 | (0.046) | all > .05 |
| M-CSF | 1.372 | (0.03) | 1.437 | (0.028) | 1.37 | (0.038) | 1.415 | (0.049) | 1.434 | (0.011) | 1.422 | (0.03) | 1.427 | (0.02) | 1.4275 | (0.021) | $ .028 |
| IL-17A | 1.452 | (0.043) | 1.515 | (0.029) | 1.284 | (0.102) | 1.316 | (0.158) | 1.504 | (0.016) | 1.494 | (0.029) | 1.472 | (0.046) | 1.472 | (0.047) | all > .05 |
| IL-1B | 1.418 | (0.035) | 1.452 | (0.058) | 1.184 | (0.228) | 1.366 | (0.093) | 1.468 | (0.041) | 1.468 | (0.045) | 1.266 | (0.038) | 1.408 | (0.048) | all > .05 |
| IL-4 | 1.09 | (0.094) | 1.3 | (0.029) | 1.166 | (0.044) | 1.126 | (0.112) | 1.254 | (0.04) | 1.28 | (0.031) | 1.408 | (0.048) | 1.266 | (0.039) | † .033 |
| IFN-g | 1.018 | (0.107) | 1.125 | (0.033) | 0.884 | (0.155) | 0.99 | (0.142) | 1.154 | (0.036) | 1.216 | (0.026) | 1.142 | (0.03) | 1.1425 | (0.031) | † .016 |
| MIP-3a | 1.442 | (0.065) | 1.487 | (0.058) | 1.424 | (0.133) | 1.315 | (0.135) | 1.578 | (0.07) | 1.536 | (0.076) | 1.43 | (0.069) | 1.43 | (0.07) | all > .05 |
| GM-CSF | 1.3 | (0.046) | 1.427 | (0.049) | 1.28 | (0.087) | 1.307 | (0.091) | 1.402 | (0.029) | 1.45 | (0.03) | 1.394 | (0.044) | 1.394 | (0.045) | † .028 |
| IL-7 | 1.298 | (0.053) | 1.33 | (0.02) | 1.29 | (0.047) | 1.037 | (0.287) | 1.394 | (0.029) | 1.432 | (0.033) | 1.417 | (0.042) | 1.4175 | (0.043) | all > .05 |
| TNFA | 1.166 | (0.069) | 1.277 | (0.037) | 1.13 | (0.076) | 1.124 | (0.098) | 1.246 | (0.014) | 1.085 | (0.106) | 1.287 | (0.054) | 1.2875 | (0.054) | all > .05 |
| VEGF | 1.45 | (0.049) | 1.507 | (0.032) | 1.5 | (0.047) | 1.304 | (0.174) | 1.56 | (0.023) | 1.55 | (0.045) | 1.438 | (0.057) | 1.438 | (0.057) | all > .05 |
| MCP-1 | 1.888 | (0.128) | 1.9 | (0.104) | 1.964 | (0.101) | 1.842 | (0.195) | 2.068 | (0.03) | 2.005 | (0.043) | 1.77 | (0.1) | 1.77 | (0.1) | all > .05 |
| IL-5 | 0.912 | (0.153) | 1.145 | (0.052) | 0.954 | (0.267) | 0.997 | (0.073) | 1.135 | (0.041) | 1.136 | (0.036) | 1.045 | (0.023) | 1.045 | (0.024) | all > .05 |
| GCSF | 0.853 | (0.203) | 1.23 | (0.045) | 1.177 | (0.013) | 1.048 | (0.12) | 1.164 | (0.021) | 1.198 | (0.012) | 1.216 | (0.039) | 1.216 | (0.04) | $* .042 $*† .025 |
| RANTES | 2.578 | (0.192) | 2.572 | (0.215) | 2.638 | (0.143) | 2.39 | (0.289) | 2.758 | (0.073) | 2.474 | (0.251) | 2.272 | (0.215) | 2.272 | (0.22) | all > .05 |
| IL-6 | 1.254 | (0.037) | 1.265 | (0.029) | 1.222 | (0.022) | 1.287 | (0.043) | 1.302 | (0.027) | 1.328 | (0.025) | 1.26 | (0.046) | 1.26 | (0.046) | all > .05 |
| GRO/KC | 1.798 | (0.097) | 1.95 | (0.01) | 1.918 | (0.039) | 1.662 | (0.214) | 1.952 | (0.028) | 1.98 | (0.008) | 1.93 | (0.012) | 1.93 | (0.012) | † .039 |
| MIP1A | 1.747 | (0.072) | 1.745 | (0.127) | 1.592 | (0.132) | 1.367 | (0.275) | 1.8 | (0.052) | 1.772 | (0.106) | 1.576 | (0.084) | 1.576 | (0.085) | * .036 |
| IL-12p70 | 1.298 | (0.074) | 1.47 | (0.043) | 1.29 | (0.128) | 1.462 | (0.011) | 1.444 | (0.025) | 1.482 | (0.021) | 1.454 | (0.024) | 1.454 | (0.024) | † .035 $ .013 |

Serum collected from pups on postnatal day 9 was analyzed using a cytokine multiplex. The data was transformed by applying the base-10 logarithm. Values represent the group mean and SEM. Asterisk, obelisk, dollar sign, dollar sign and asterisks, dollar sign, asterisk and obelisk, indicate a significant main effect of pump, fluid, sex, sex by pump, sex by pump by fluid respectively.

| Supplementary Results Table ii: Cytokine expression in pups’ serum at postnatal day 21 | | | | | | | | | | | | | | | | | |
| --- | --- | --- | --- | --- | --- | --- | --- | --- | --- | --- | --- | --- | --- | --- | --- | --- | --- |
|  | Sham/Veh Mean (SEM) | | | | Met/Veh Mean (SEM) | | | | Sham/NaB Mean (SEM) | | | | Met/NaB Mean (SEM) | | | |  |
|  | Male (n=4) | | Females (n=6) | | Male (n=4) | | Females (n=6) | | Male (n=6) | | Females (n=5) | | Male (n=3) | | Females (n=5) | | *p-*value |
| IL-13 | 2.587 | (0.081) | 2.701 | (0.081) | 2.302 | (0.203) | 2.422 | (0.212) | 2.56 | (0.14) | 2.788 | (0.03) | 2.59 | (0.109) | 2.616 | (0.142) | all > .05 |
| IL-10 | 2.285 | (0.023) | 2.305 | (0.026) | 2.257 | (0.074) | 2.328 | (0.055) | 2.261 | (0.04) | 2.32 | (0.029) | 2.23 | (0.03) | 2.275 | (0.029) | all > .05 |
| IL-1A | 2.642 | (0.017) | 2.6 | (0.017) | 2.652 | (0.095) | 2.371 | (0.405) | 2.676 | (0.035) | 2.652 | (0.045) | 2.63 | (0.063) | 2.66 | (0.046) | all > .05 |
| IL-2 | 3.427 | (0.048) | 3.365 | (0.008) | 3.395 | (0.094) | 3.375 | (0.064) | 3.335 | (0.039) | 3.427 | (0.031) | 3.313 | (0.037) | 3.375 | (0.089) | all > .05 |
| M-CSF | 1.225 | (0.285) | 1.135 | (0.095) | 1.123 | (0.178) | 1.503 | (0.25) | 1.158 | (0.073) | 1.192 | (0.026) | 1.295 | (0.215) | 1.053 | (0.354) | all > .05 |
| IL-1B | 2.037 | (0.284) | 1.816 | (0.157) | 1.842 | (0.08) | 1.45 | (0.456) | 1.886 | (0.112) | 2.172 | (0.076) | 1.96 | (0.348) | 1.797 | (0.134) | all > .05 |
| IL-4 | 2.792 | (0.041) | 2.843 | (0.021) | 2.682 | (0.104) | 2.832 | (0.074) | 2.781 | (0.044) | 2.814 | (0.03) | 2.73 | (0.075) | 2.828 | (0.051) | all > .05 |
| IFN-g | 2.452 | (0.137) | 2.516 | (0.09) | 2.227 | (0.203) | 2.395 | (0.155) | 2.47 | (0.103) | 2.608 | (0.033) | 2.386 | (0.154) | 2.48 | (0.115) | all > .05 |
| MIP-3a | 1.527 | (0.049) | 1.565 | (0.025) | 1.6 | (0.125) | 1.378 | (0.197) | 1.673 | (0.031) | 1.747 | (0.106) | 1.73 | (0.116) | 1.672 | (0.008) | † .035 |
| GM-CSF | 2.272 | (0.261) | 2.138 | (0.133) | 2.152 | (0.121) | 2.278 | (0.174) | 2.16 | (0.113) | 2.19 | (0.045) | 2.213 | (0.329) | 2.097 | (0.211) | all > .05 |
| IL-7 | 2.242 | (0.27) | 2.088 | (0.096) | 2.112 | (0.069) | 1.902 | (0.306) | 2.091 | (0.084) | 2.157 | (0.043) | 2.146 | (0.305) | 2.112 | (0.153) | all > .05 |
| TNFA | 2.667 | (0.083) | 2.515 | (0.227) | 2.367 | (0.324) | 2.59 | (0.081) | 2.693 | (0.054) | 2.82 | (0.036) | 2.726 | (0.049) | 2.692 | (0.122) | all > .05 |
| VEGF | 2.16 | (0.125) | 2.268 | (0.125) | 2.175 | (0.149) | 2.18 | (0.092) | 2.125 | (0.055) | 2.182 | (0.012) | 2.136 | (0.157) | 2.17 | (0.094) | all > .05 |
| MCP-1 | 3.222 | (0.095) | 3.144 | (0.048) | 3.188 | (0.064) | 3.206 | (0.051) | 3.196 | (0.031) | 3.177 | (0.251) | 3.2 | (0.113) | 3.27 | (0.13) | all > .05 |
| IL-5 | 2.852 | (0.006) | 2.872 | (0.006) | 2.83 | (0.03) | 2.93 | (0.042) | 2.881 | (0.027) | 2.884 | (0.025) | 2.823 | (0.024) | 2.862 | (0.021) | all > .05 |
| RANTES | 2.945 | (0.096) | 3.061 | (0.07) | 3.062 | (0.041) | 3.056 | (0.051) | 3.023 | (0.061) | 3.02 | (0.008) | 2.963 | (0.065) | 2.935 | (0.091) | all > .05 |
| IL-6 | 2.513 | (0.131) | 2.692 | (0.067) | 2.433 | (0.135) | 2.697 | (0.154) | 2.637 | (0.072) | 2.672 | (0.106) | 2.475 | (0.265) | 2.542 | (0.13) | $ .003 |
| GRO/KC | 2.575 | (0.105) | 2.512 | (0.085) | 2.496 | (0.049) | 2.468 | (0.062) | 2.476 | (0.031) | 2.507 | (0.021) | 2.406 | (0.232) | 2.592 | (0.187) | all > .05 |
| IL-17A | 1.572 | (0.034) | 1.65 | (0.025) | 1.484 | (0.087) | 1.632 | (0.07) | 1.638 | (0.039) | 1.64 | (0) | 1.566 | (0.013) | 1.627 | (0.04) | all > .05 |
| MIP1A | 1.917 | (0.205) | 1.846 | (0.086) | 1.825 | (0.059) | 1.825 | (0.082) | 1.868 | (0.075) | 1.89 | (0) | 1.893 | (0.224) | 1.86 | (0.132) | all > .05 |
| IL-12p70 | 2.49 | (0.016) | 2.598 | (0.024) | 2.444 | (0.117) | 2.525 | (0.057) | 2.518 | (0.067) | 2.62 | (0) | 2.443 | (0.094) | 2.522 | (0.048) | $ .039 |

Serum collected from pups on postnatal day 21 was analyzed using a cytokine multiplex. The data was transformed by applying the base-10 logarithm. Each value represents the group mean and SEM. Obelisk and dollar sign indicate a significant main effect of fluid or sex respectively.

Supplementary Results Table iii: Sholl and Skeleton analysis of microglia in the dentate gyrus at P21

|  | Sham/Veh Mean (SEM) | | | | Met/Veh Mean (SEM) | | | | Sham/NaB Mean (SEM) | | | | Met/NaB Mean (SEM) | | | |
| --- | --- | --- | --- | --- | --- | --- | --- | --- | --- | --- | --- | --- | --- | --- | --- | --- |
|  | Male (n=6) | | Females (n=4) | | Male (n=6) | | Females (n=6) | | Male (n=5) | | Females (n=3) | | Male (n=5) | | Females (n=4) | |
| Intersecting radii | 9.5 | (1.602) | 4.143 | (0.875) | 7.690 | (1.319) | 7.762 | (1.420) | 6.4 | (0.921) | 5.810 | (1.061) | 6.524 | (1.206) | 6.071 | (1.028) |
| Max intersections | 4.714 | (0.382) | 4.143 | (0.309) | 3.452 | (0.305) | 4.262 | (0.275) | 3.652 | (0.561) | 4.286 | (0.595) | 4.095 | (0.266) | 3.821 | (0.380) |
| Ramification index | 1,889 | (0.178) | 1.467 | (0.177) | 1.538 | (0.096) | 1.790 | (0.124) | 1.466 | (0.191) | 1.774 | (0.344) | 1.397 | (0.079) | 1.432 | (0.146) |
| Whole image cell count | 18.333 | (2.404) | 14.400 | (1.631) | 15.333 | (1.706) | 16.333 | (3.528) | 13.600 | (1.860) | 13.00 | (2.082) | 15.200 | (1.772) | 12.250 | (1.315) |
| Whole image branches | 17.987 | (2.026) | 20.600 | (3.862) | 15.875 | (0.611) | 16.593 | (1.507) | 13.636 | (2.213) | 21.864 | (4.258) | 15.527 | (1.719) | 19.131 | (3.880) |
| Whole image junctions | 8.241 | (0.986) | 9.474 | (1.935) | 7.277 | (0.322) | 7.594 | (0.768) | 6.054 | (1.084) | 10.248 | (2.175) | 7.039 | (0.877) | 8.964 | (1.928) |
| Whole image end point voxels | 10.166 | (0.908) | 10.983 | (1.559) | 8.826 | (0.325) | 9.331 | (0.708) | 8.113 | (1.079) | 11.912 | (1.877) | 8.823 | (0.728) | 10.186 | (1.646) |
| Whole image junction voxels | 28.779 | (3.330) | 33.774 | (7.240) | 24.651 | (0.835) | 26.073 | (2.504) | 21.491 | (3.481) | 34.091 | (7.267) | 24.554 | (2.879) | 30.271 | (6.240) |
| Whole image slab voxels | 145.150 | (18.823) | 144.386 | (27.964) | 122.238 | (8.805) | 122.801 | (12.238) | 110.048 | (23.578) | 191.535 | (56.410) | 117.885 | (14.351) | 145.487 | (33.874) |
| Whole image average branch length | 3.873 | (0.270) | 3.486 | (0.253) | 3.644 | (0.258) | 3.594 | (0.244) | 3.760 | (0.209) | 3.959 | (0.403) | 3.608 | (0.098) | 3.649 | (0.164) |
| Whole image triple points | 7.211 | (0.826) | 7.899 | (1.600) | 6.225 | (0.269) | 6.661 | (0.757) | 5.164 | (1.001) | 9.234 | (2.141) | 6.063 | (0.795) | 7.902 | (1.575) |
| Whole image quadruple points | 0.904 | (0.204) | 1.358 | (0.328) | 0.980 | (0.087) | 0.803 | (0.056) | 0.797 | (0.116) | 0.882 | (0.115) | 0.839 | (0.093) | 0.967 | (0.373) |
| Whole image maximum branch length | 10.541 | (0.965) | 8.535 | (0.824) | 9.555 | (0.845) | 9.503 | (0.885) | 9.694 | (1.345) | 12.286 | (2.382) | 9.265 | (0.340) | 10.189 | (1.128) |
| Whole image longest shortest path | 35.111 | (3.932) | 31.617 | (4.385) | 30.629 | (2.082) | 31.091 | (2.532) | 28.526 | (5.258) | 42.822 | (9.320) | 29.724 | (2.971) | 34.211 | (5.207) |
| Individual cells branches | 21.095 | (2.941) | 20.535 | (4.586) | 13.942 | (3.144) | 22.190 | (6.195) | 14 | (2.267) | 12.828 | (1.571) | 18.971 | (3.088) | 18.143 | (3.111) |
| Individual cells junctions | 9.786 | (1.439) | 9.429 | (2.225) | 6.2 | (1.448) | 10.476 | (3.268) | 6.171 | (1.087) | 5.771 | (0.806) | 8.743 | (1.537) | 8.428 | (1.569) |
| Individual cells end point voxels | 11.643 | (1.288) | 11.464 | (2.146) | 8.286 | (1.643) | 11.762 | (2.436) | 8.543 | (1.039) | 8.343 | (0.804) | 10.8 | (1.579) | 10.035 | (1.309) |
| Individual cells junction voxels | 32.357 | (4.188) | 32.321 | (8.093) | 24.657 | (4.773) | 35.714 | (10.655) | 21.857 | (4.022) | 22.371 | (3.780) | 29.485 | (5.114) | 28.464 | (4.665) |
| Individual cells slab voxels | 203.786 | (18.500) | 182.179 | (31.553) | 150.114 | (30.943) | 244.000 | (66.337) | 143.000 | (27.454) | 136.816 | (19.451) | 182.714 | (33.173) | 185.250 | (29.712) |
| Individual cells average branch length | 4.749 | (0.284) | 4.425 | (0.265) | 5.764 | (0.559) | 5.081 | (0.283) | 4.989 | (0.207) | 5.390 | (0.533) | 4.889 | (0.224) | 5.127 | (0.224) |
| Individual cells triple points | 8.595 | (1.164) | 8.143 | (1.864) | 4.971 | (1.239) | 9.381 | (3.126) | 5.400 | (0.891) | 4.800 | (0.746) | 7.886 | (1.551) | 7.500 | (1.397) |
| Individual cells quadruple points | 1.190 | (0.339) | 1.143 | (0.408) | 0.943 | (0.318) | 1.000 | (0.218) | 0.657 | (0.184) | 1.028 | (0.223) | 0.800 | (0.132) | 0.893 | (0.327) |
| Individual cells maximum branch length | 13.952 | (1.170) | 11.750 | (0.890) | 15.109 | (1.261) | 15.390 | (2.314) | 13.834 | (1.938) | 14.464 | (0.80) | 13.783 | (0.896) | 15.148 | (0.606) |
| Individual cells longest shortest path | 48.219 | (2.518) | 38.822 | (3.904) | 38.469 | (5.360) | 52.182 | (9.580) | 37.237 | (6.330) | 37.910 | (4.130) | 41.875 | (5.830) | 43.653 | (4.509) |
| Individual cells cell body area | 53.164 | (9.588) | 98.872 | (23.552) | 52.681 | (5.961) | 60.222 | (8.575) | 69.950 | (8.684) | 68.397 | (8.133) | 60.194 | (6.141) | 44.537 | (17.824) |
| Individual cells cell body perimeter | 32.057 | (1.953) | 38.977 | (3.453) | 28.770 | (1.278) | 30.286 | (2.383) | 32.766 | (2.011) | 33.064 | (2.880) | 30.367 | (1.746) | 23.558 | (8.023) |

Each value represents the group mean and SEM.
